# Maximum-likelihood model fitting for quantitative analysis of SMLM data

**DOI:** 10.1101/2021.08.30.456756

**Authors:** Yu-Le Wu, Philipp Hoess, Aline Tschanz, Ulf Matti, Markus Mund, Jonas Ries

## Abstract

Quantitative analysis is an important part of any single-molecule localization microscopy (SMLM) data analysis workflow to extract biological insights from the coordinates of the single fluorophores, but current approaches are restricted to simple geometries or do not work on heterogenous structures.

Here, we present LocMoFit (**Loc**alization **Mo**del **Fit**), an open-source framework to fit an arbitrary model directly to the localization coordinates in SMLM data. Using maximum likelihood estimation, this tool extracts the most likely parameters for a given model that best describe the data, and can select the most likely model from alternative models. We demonstrate the versatility of LocMoFit by measuring precise dimensions of the nuclear pore complex and microtubules. We also use LocMoFit to assemble static and dynamic multi-color protein density maps from thousands of snapshots. In case an underlying geometry cannot be postulated, LocMoFit can perform single-particle averaging of super-resolution structures without any assumption about geometry or symmetry. We provide extensive simulation and visualization routines to validate the robustness of LocMoFit and tutorials based on example data to enable any user to increase the information content they can extract from their SMLM data.

## Introduction

Single-molecule localization microscopy (SMLM), such as PALM (photoactivated localization microscopy^1,2^) or STORM (stochastic optical reconstruction microscopy^3^), enables nanometer optical super-resolution and has widespread applications in cell and structural biology. SMLM is based on precisely localizing single switchable fluorophores that are sparsely activated in individual camera frames. From the coordinates of the fluorophores, determined in thousands of frames, a super-resolution image is reconstructed. Visual inspection of this image can provide insights into the investigated biological system, but to gain reliable mechanistic insights, a quantitative analysis is indispensable. This is especially true when large amounts of data are created using high-throughput SMLM^4–7^, which precludes a visual analysis. Such a quantitative analysis aims at informing on properties of the biological system or at probing functional differences among different conditions with statistical confidence.

Standard image analysis algorithms can be applied directly to a rendered pixelated superresolution image, but often their performance is limited due to the very different image formation process and information content in SMLM compared to standard fluorescence microscopy. The primary data in SMLM, but also in the new MINFLUX^8^ super-resolution technology, are not the reconstructed images, but a list of coordinates of fluorophores, often with additional information such as an estimate of the localization uncertainty. Thus, algorithms that directly use these coordinates can exploit this additional information and can produce more accurate and robust results^9^. These algorithms can be assigned to several classes^9^: *Spatial descriptive statistics*, such as pair-correlation^10^ or Ripley’s K-function^11^, inform on the clustering state and cluster sizes in specific regions of interest. After segmentation of individual structures, they can be *classified*^12^, or analyzed with a *geometric analysis*. Examples include size estimation from the standard deviation of the coordinates, fitting of single or double Gaussians to line profiles^13–15^, or fitting of a circle to extract the diameter of ring-shaped structures^16,17^. However, to date, only simple models have been used that cannot adequately describe the complex geometry of many cellular structures. In *particle averaging* or fusion, an approach adapted from electron microscopy, hundreds of particles or structures are registered and averaged to result in a final model with improved resolution and signal^18–20^. However, this approach relies on the underlying biological structures to be identical with minimal biological variability.

Neither of these approaches is applicable to the most typical scenario of SMLM data analysis: Usually, some aspects of the geometry underlying the structure of interest can be guessed from visual inspection of the super-resolution images or from prior knowledge based on structural biology techniques. The data analysis task then consists of first selecting the most likely geometry from a class of possible models and second extracting precise parameters describing this geometry. Such analysis would be applicable to individual structures and thus could quantify biological and functional heterogeneities. However, to our knowledge, algorithms for such analysis of SMLM data do not yet exist.

Here, we developed **Loc**alization **Mo**del **Fit** (LocMoFit; Fig. 1), a general framework to fit an arbitrary model to coordinate-based SMLM data. Based on maximum likelihood estimation (MLE), it identifies the most likely model from a class of models and estimates the most likely parameters of the model that describe the experimental structure. If the underlying geometry cannot be guessed, LocMoFit can be used for particle averaging to calculate an average model under the assumption of identical structures. Advanced visualization routines and a simulation engine allow for efficient validation and quality control. LocMoFit, written in MATLAB, has an API for integration into own code and can be easily extended by user-defined models. Seamless integration of LocMoFit in the open-source analysis platform SMAP^21^ provides access to plenty of tools for SMLM for localization, post-processing and quantification. Distributed as open source and with numerous examples and extensive documentation, LocMoFit will enable many researchers to perform quantitative analysis of their data with unprecedented efficiency, accuracy and statistical power.

**Figure 1.**
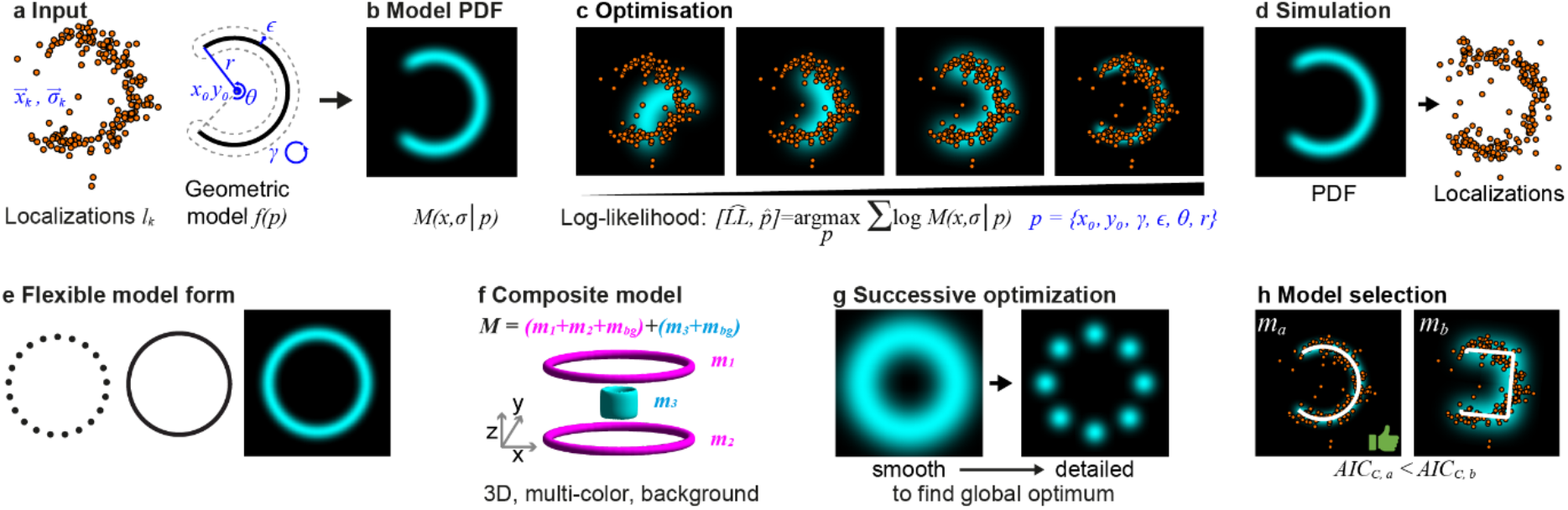
Overview of LocMoFit. **a-c, Workflow** of the fitting procedure. **a**, Inputs of LocMoFit are the spatial coordinate 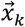 and localization precision 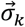 of each localization *k* and a geometric model *f* parameterized by parameters *p*. **b**, First the probability density function (PDF) 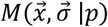 of the input model is constructed. **c,** From the model PDF the likelihood is calculated that the model describes the data. A maximum likelihood estimation (MLE) routine searches in the parameter space and maximizes the log-likelihood to find parameter values that best describe the localizations. In the example, a 2D arc model (cyan), parameterized by positions *x*_0_, *y*_0_, rotational angle *γ*, linkage error *ϵ*, arc opening angle *θ*, and radius *r*, is fitted to the single-color data (orange dots). **d-h, Features of the framework. d, Simulation engine** for validation. Labels are simulated as samples drawn from the PDF and localizations are then calculated based on fluorophore properties including photon counts, re-blinks, and labeling efficiency. **e**, The framework supports **flexible model forms** including discrete/continuous models and images. **f**, It can assemble **complex composite models** from simple ones and supports 3D and multi-color data. In the example, the composite model *M* is formed by combining two ring models (*m*_1_ and *m*_2_) and one cylindrical model (*m*_3_), which are assigned to different channels, represented by different colors. Background models *m_bg_* are incorporated channel-wise. **g**, LocMoFit allows **multi-step successive optimization** to avoid local optima. In the example, a smooth, continuous ring model is used to robustly estimate approximate parameters. These are then passed on as start parameters for a fit with a detailed eight-fold symmetric model with discrete corners. **h**, **Model selection**. Based on the corrected version of Akaike information criterion (AIC_C_) reported by LocMoFit, the model that best describes the data can be selected. In the example, the arc model *m_a_* has a smaller AIC_C_ than the bucket model *m_b_*, reflecting that it is a better model for describing the example localizations.

## Results

### Localization Model Fit (LocMoFit)

LocMoFit fits a geometric model *f*(*p*) to a set of *K* localizations 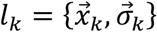 (Fig. 1a-c) in a region with a defined boundary that we call ‘site’, which corresponds to one biological structure or ‘particle’. *l_k_* are obtained by fitting camera images with a model of the point spread function and are described by their coordinates 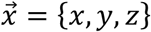, the uncertainties of the coordinates 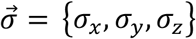 for 3D data and 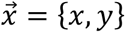 and 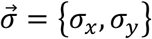 for 2D data. We demonstrate the workflow using an arc site generated using the simulation functionality of LocMoFit (Fig. 1d). *f*(*p*) describes the spatial distribution of the imaged fluorophores and is parameterized by the set of parameters *p*. Our approach is to use maximum likelihood estimation (MLE) to find the set of parameters 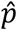 that, together with *f*(*p*), best describes the measured *l_k_* (Fig. 1c). For this, we first use *f*(*p*) to calculate the probability density function (PDF) 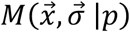 that describes the probability that, if we acquire a single localization *l* with the uncertainty 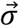 at random, it is found at the coordinate 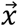. The likelihood to obtain the set *l_k_* of *K* localizations in a measurement is then given by the product of individual probabilities:

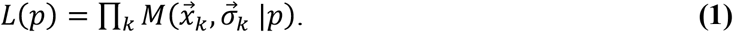

We then use an optimization algorithm to find the parameters 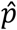 that maximize *L*(*p*):

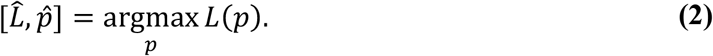

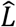 denotes the estimate of the maximum likelihood. For efficiency and to prevent a small joint probability from being rounded to zero, the natural logarithm of the likelihood, the log-likelihood *LL*(*p*), is usually used in practice (see Fig. 1c).

The PDF *M* is constructed from the geometric model *f*(*p*). *f*(*p*) is defined either in a continuous or a discrete form, or supplied as an image (Fig. 1e). A continuous *f*(*p*) describes the shapes formed by the fluorophores such as one-dimensional lines (e.g. filaments or rings) or two-dimensional surfaces (spheres, patches), while a discrete *f*(*p*) describes the exact fluorophore positions.

In the simplest case, 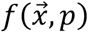 directly describes whether a fluorophore can be found at the position 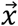. Due to a limited localization precision 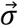, the position of the localization coordinate is not equal to the true position of the fluorophore, but is instead randomly displaced by 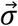. This localization uncertainty is used when constructing the PDF by convolving 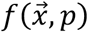 with a Gaussian function with standard deviations given by the mean of the localization precision 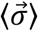 (⊗ denotes the convolution):

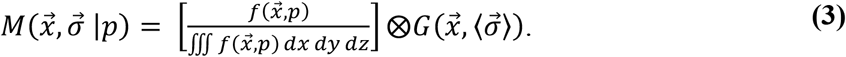

Alternatively, LocMoFit can take into account that each localization has its specific lateral and axial localization uncertainties. In this scenario, the model 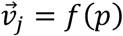 directly specifies the expected coordinates 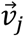 of the in total *J* fluorophore positions in the model. The likelihood that the localization 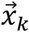 stems from the fluorophore 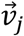 is described by a Gaussian function and depends on the distance between 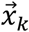 and 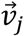 and the localization precision 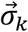. To construct the model *M* for this single localization, we sum over all model localizations *j*:

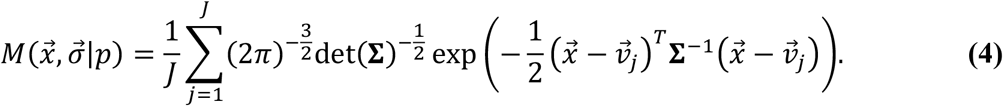

The parameters *p* consist of intrinsic parameters *p^i^* that directly determine the shape of the model and extrinsic parameters *p^e^* that describe the rigid transformation and rescaling of the model. 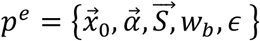 includes the position of the model 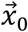, the orientation, described by the rotation angles 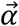 around the coordinate axes, an optional global scaling factor 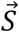, and the proportional weight *w_b_* of a constant background PDF *M_bg_* that accommodates the localizations that cannot be described by the geometric PDF. An optional additional uncertainty *ϵ* accommodates an uncertainty that cannot be described by the localization precision, such as a linkage error of the fluorophore (e.g., due to immunolabeling with primary and secondary antibodies), small-scale deformations of the structure that are not described by the model or residual instabilities of the microscope. 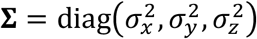 is the diagonal matrix of the square of the localization uncertainties and det(**Σ**) is the determinant of the matrix **Σ**. When *p^e^* is fitted during the MLE, the expected coordinates 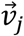 are replaced by 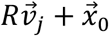, the uncertainty 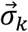 by 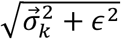, and the geometric PDF 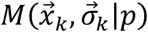 by 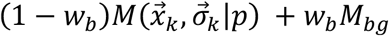. From the optimization, we obtain the parameter estimates 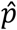 along with the 95% confidence intervals of each fit parameter.

To describe a more complex geometry, a composite model (magenta only in Fig. 1f) PDF *M_c_* can be formed by a linear combination of sub models *M_m_* that share the same background:

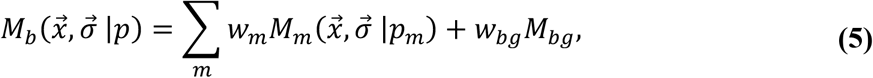

where the sum of weights ∑*_m_w_m_* + *w_b_* = 1 for normalization and *p* = {*p_m_,m* = 1…*N*} for in total *N* component models.

When fitting a composite model to more than one color at a time (e.g. both colors in Fig. 1e), the model PDF can be constructed as

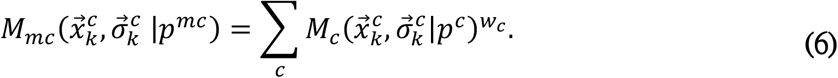

Here, each localization is not only described by its coordinate and localization precision, but also its color *c*. Note that each single-color PDF 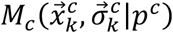 is evaluated only with the localizations of the corresponding color. *w_c_* is the weight for each color and is by default set to 1 and therefore not active. If the colors have very different numbers of localizations, the fit can be dominated by one color. In this case, one can set *w_c_* = ∑*_c_ K_c_*/*K_c_*, the inverse of the fraction of localizations with the respective color, to scale to the number of multiplications in Equation (1) and thus give each color the same weight in the fit. Equation (6) is the general form of the model PDF, which can describe a vast class of biological structures.

To prevent the optimization from getting stuck in a local maximum of the likelihood, LocMoFit allows the user to chain several fitting steps with different models (usually in the order from smooth to detailed), passing on the parameter estimates from the previous step to the next one as the initial parameters (Fig. 1g).

Since the likelihood itself is a measure of the goodness of fit, the model that best describes the data can be identified by comparing the log-likelihood, or more precisely the Akaike information criterion^22^, of different models fitted to the same data (Fig. 1h).

The probabilistic likelihood *L*(*p*) used in LocMoFit is related to cross-correlation used in other studies for measuring similarity between sites^23,24^. Indeed, we can obtain the cross-correlation cost function by replacing the multiplication of *L*(*p*) (see equation (1)) with a summation:

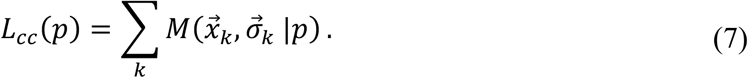

This cost function corresponds to a general cross-correlation cost function^23,24^ under the assumption of a constant localization uncertainty for the template. Also, it is closely related to the Bhattacharya cost function that was previously used for particle fusion in SMLM^18,19^.

### Simulation and validation

To test the robustness of specific model fitting pipelines, we included a comprehensive simulation engine in LocMoFit (Fig. 1d), as an extension of the engine applied in our previous work^17^. This engine uses a realistic description of fluorophore blinking (see **Methods**) to generate synthetic localization data from any model. Fitting these synthetic structures allows comparing the fitted model parameters to the ground truth. We usually validate a fitting pipeline with simulations before applying it to experimental data.

We systematically investigated on the example of the nuclear pore complex (NPC) how the precision and accuracy of the fit parameters depends on localization precision, background localizations, labeling efficiency and number of localizations per fluorophore (re-blinks) (see Extended Data Figure 1 and Extended Data Figure 2 and **Methods**).

As Nup96 has been assigned into EM-densities^25^, we know that it distributes in two rings, exhibits an 8-fold rotational symmetry in the NPC and that each symmetric unit is occupied by two copies per ring. Based on this prior knowledge, we constructed our detailed model of the NPC (Extended Data Figure 1a). We simulated localization data of NPCs with pre-defined parameters, the ‘ground truth’, as shown in Supplementary Table 1. We acquired the parameter estimates by fitting the simulated data with the model and computed the errors of the estimates. As shown in Extended Data Figure 2, the errors in the estimated parameters (e.g., position, rotation, ring radius or distance) are close to zero with small spreads, showing that the fitting is unbiased and reliable for a large range of experimental conditions. Too low labeling efficiencies, however, resulted in a bias of the average ring separation to smaller distances (Extended Data Figure 2a). The reason is that under these labeling efficiencies a subset of the NPCs has one ring entirely unlabeled, and the remaining ring is well fitted by the two-ring model with a small separation parameter (Extended Data Figure 3). This highlights the importance of simulations for identifying potential factors to be considered when interpreting results. As expected, the precision in the parameter estimates was the highest for brightest fluorophores, and lowest backgrounds. The remaining fitting parameters showed negligible systematic errors.

### Extracting structural parameters from individual sites

LocMoFit allows determining specific and meaningful structural parameters from individual sites without any averaging, which can be used to gain structural insights into multi-protein assemblies and to investigate biological heterogeneity. Here, we demonstrate the power and flexibility of LocMoFit on two biological structures that have been used extensively as test structures in SMLM, the nuclear pore complex (NPC) and microtubules.

We imaged the protein Nup96 tagged with SNAP-tag in the NPC in a genome edited cell line^17,26^ and obtained hundreds of NPC structures per field of view (Fig. 2a-f). In 3D reconstructions, Nup96 appeared as two parallel rings (Fig. 2a,b). We thus fitted the NPC with a two-ring model to extract the approximate positions and orientations of each site as initial parameters for a more detailed model fit.

**Figure 2.**
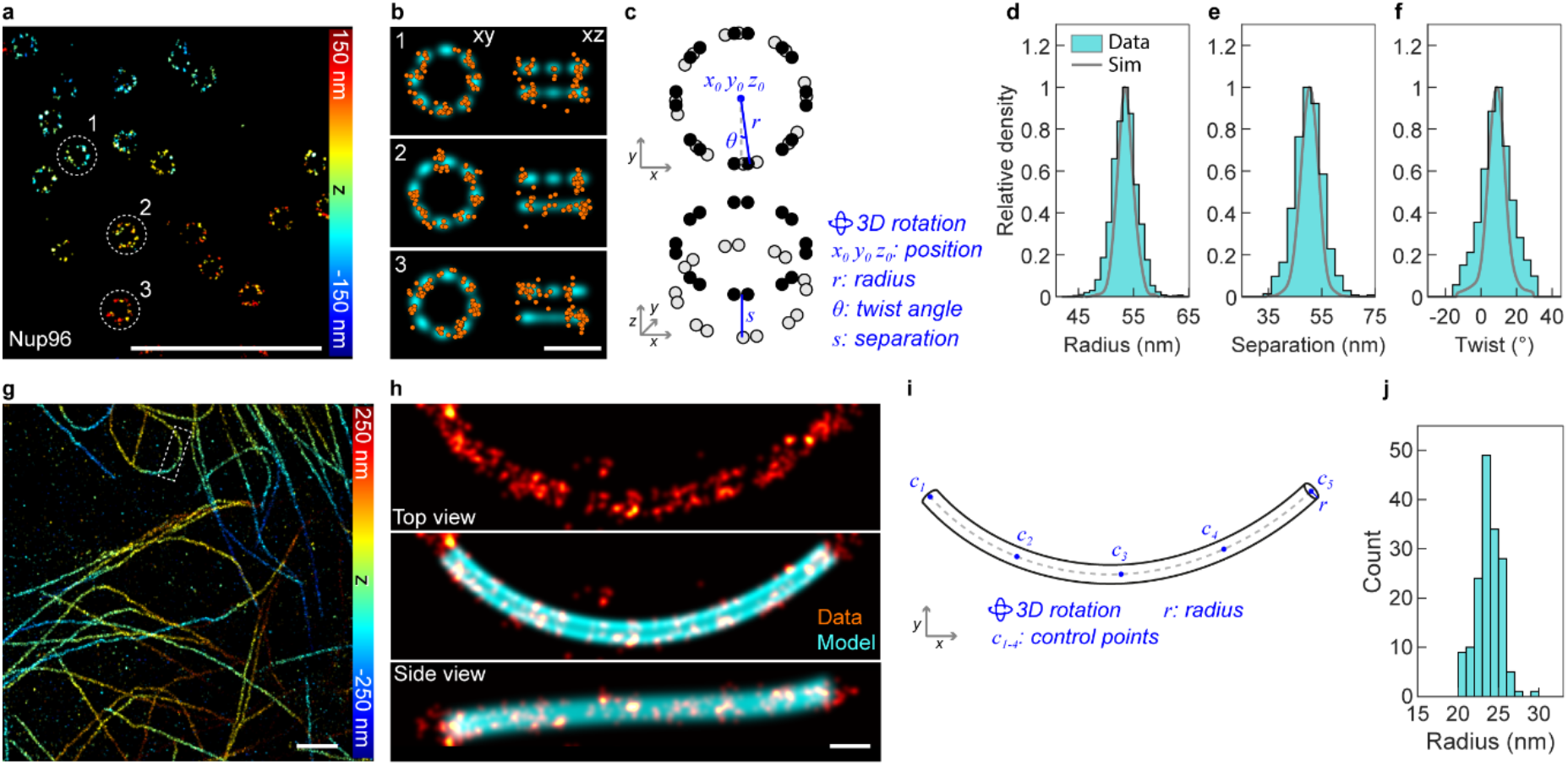
Quantifying structures at single sites. **a-f**, **Quantification of NPCs**. **a**, **Representative image** of the Nup96-labeled NPCs (Nup96-SNAP-AF647) in a 3D dataset (top view). **b**, single NPCs (localizations in orange) as indicated in **a** are fitted with the eight-fold symmetric model (cyan) shown in (**c**). The model is parameterized by the listed parameters (blue). **d-e**, Histograms of three fitted parameters: radius *r* = 53.4 ± 2.3 nm (**d**), separation *s* = 50.2 ± 5.8 nm (**e**), and twist *θ* = 8.8 ± 9.0° (**f**). ‘Sim’ stands for simulation (gray). Sample size: *n*_s_ = 3,524, *n*_c_ = 5. **g-j**, **Quantification of microtubules**. **g**, Representative image of immunolabeled microtubules in a 3D dataset (top view). **h**, One microtubule segment (red) as indicated in **g** is fitted by the linear-tube model (**i**). The fitted model is indicated in cyan. **i**, The linear-tube model parameterized by the listed parameters (blue), the control points *c_i_* define a cubic spline. **j**, Histogram of the fitted radius *r* = 24.1 ± 3.4 nm, based on segments of 1 μm length. Sample size *n*_s_ = 161, *n*_c_ = 1. Shown values are mean± s.d., based on *n*_s_ sites in total *n*_c_ cells. Scale bars, 1 μm (**a,g**), 100 nm (**b,h**).

We fitted each nuclear pore with the NPC model (as shown in Fig. 2c, the same as Extended Data Figure 1a) to extract structural parameters of NPCs: the radius *r* of the rings, the separation s between the rings and the azimuthal ‘twist’ angle *θ* between the rings. For the mean values ± standard deviations we found *r* = 53.4 ± 2.3 nm and *θ* = 8.8 ± 9.0°, in line with previously reported values^17^. As depth-dependent aberrations lead to a scaling in *z* depending on the position *z*_0_ of the NPC above the coverslip^27^, we used the measured s to directly calibrate and correct for this distortion (see **Methods** and Extended Data Figure 4). To investigate if the variation of the parameters is technical or due to biological heterogeneities, we compared our results to simulations (Extended Data Figure 5) and indeed found a larger spread of the experimental parameters (Fig. 2d-f, green compared to gray), hinting to sources of variations that we did not consider in the model, possibly biological heterogeneity. These results demonstrate that LocMoFit allows investigating the distribution of parameters for heterogenous structures.

Next, we demonstrated analysis of extended structures with LocMoFit on the example of immunolabeled microtubules (Fig. 2g-j, original data from Speiser et al.^28^). The apparent diameter of the ring is of interest, because it directly informs on the linkage error induced by the primary and secondary antibodies^29^ in indirect immunolabeling. As microtubules are generally curved, in the past the radius was usually measured only on short segments (less than 500 nm long) using a geometric fit to the cross-sectional profile^13,29^, risking a bias from low labeling densities and residual curvature. In LocMoFit, we implemented a fitting model that describes a curved tube (Fig. 2i) and thus can trace extended (micrometer-long) curved microtubule segments (Fig. 2h). We measured the apparent radius *r* of the immunolabeled microtubules as 24.1 ± 3.4 nm, 11.6 nm larger than the true mean outer radius (12.5 nm) of microtubules themselves, and similar to the reported mean radius of the shell of fluorophores in indirectly immunolabeled microtubles^29^.

### Model Selection

Using a model that faithfully approximates the biological structure is key to performing a meaningful analysis in LocMoFit. We can use LocMoFit to select the best out of a class of models by comparing the corrected version of Akaike information criterion^22^ (AIC_C_) after fitting. AIC_C_ is a derivation from maximum likelihood, with a penalty for the number of free parameters *P* and with a correction for sample size, here the number of localizations *K* (see **Methods**)^22^: *AIC_C_* = *AIC* + (2*P*^2^ + 2*P*)/(*K* – *P* – 1), where 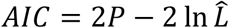. 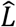 is the maximum likelihood acquired by Equation (2). For visualization, we additionally normalize the *AIC_C_* by the number of localizations. In practice, we would like to choose a model with fewer parameters and yet a larger maximum likelihood. Therefore, the smallest AIC indicates the best model when fitting the same data. To validate this idea, we fitted different models to each NPC in the Nup96 dataset (Fig. 2a,b). These models were rotationally symmetric with different symmetries (from 6-fold to 10-fold, see insets in Fig. 3a). The 8-fold symmetric model clearly shows the lowest AIC_C_ overall, in line with the known symmetry of the NPC^25^. To further validate the model-selection functionality of LocMoFit, we used its simulation engine to generate NPCs with different rotational symmetries. We demonstrate that the cumulative distributions allowed identifying the correct symmetry as a matching symmetry always had the lowest AIC_C_ (Extended Data Figure 6a). The differences in AIC_C_ decreased with increasing similarity of the models, rendering the distinction of a 10-fold symmetric model from a 9-fold pore more challenging (Extended Data Figure 6a). On the single-site level, a proper identification of the correct model is not always possible due to the relatively large variance of the AIC_C_ (Fig. 3b). Even in the simulations, a small but noticeable fraction of 8-fold symmetric NPCs have a lower AIC_C_ when fitted with a 6-fold symmetric model than an 8-fold symmetric model (Extended Data Figure 6b). A different symmetry (e.g., the 6-fold from the 8-fold), if present, stands out only when it has a comparably large population, as shown by the simulations in Extended Data Figure 6c.

**Figure 3.**
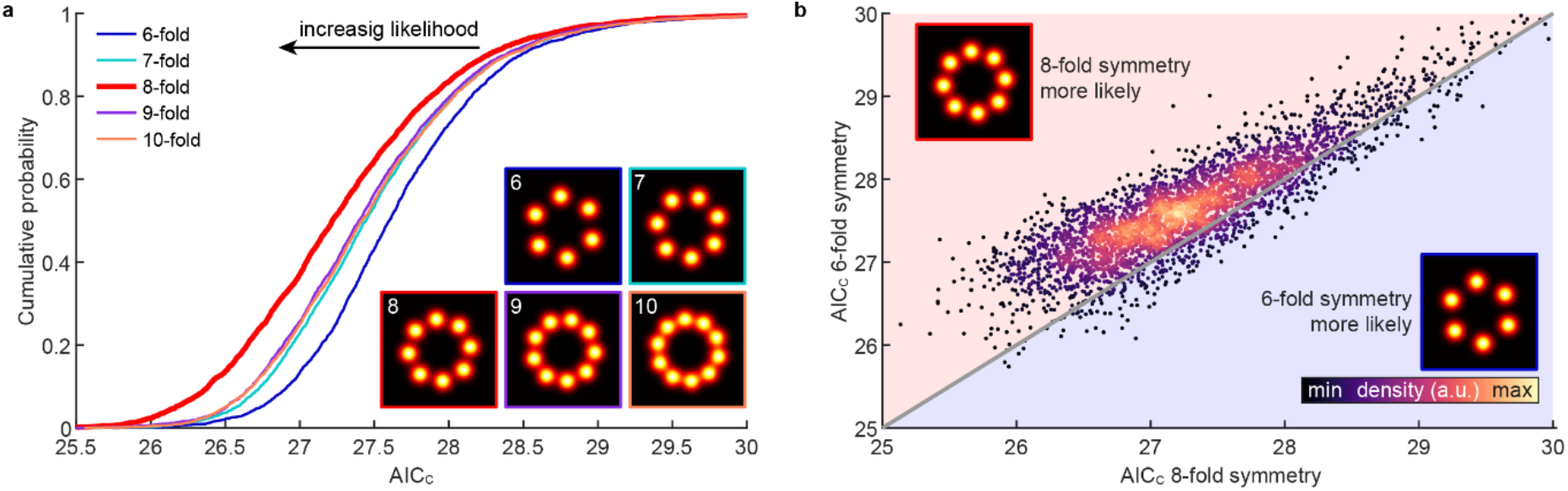
Model selection. **a**, Cumulative distribution of the normalized AIC_C_ acquired by fitting the same experimental NPC dataset with models having different rotational symmetries. The AIC_C_ was normalized by the number of localizations and assumes lower values for better fits. The (correct) 8-fold symmetric model corresponds to the lowest normalized AIC_C_. **b**, Scatter plot showing AIC_C_ of fitting the same sites with models having 6-fold and 8-fold rotational symmetries. The gray diagonal line indicates equal AIC_C_. Sample size: *n*_s_ = 3,524, *n*_c_ = 5.

### Static and dynamic protein density maps

Multi-color microscopy is widely used for studying multi-protein assemblies^6,20,30,31^. However, especially in SMLM, the number of simultaneous labels in the same sample is still a bottleneck because of spectral overlap and different optimal imaging conditions for different fluorophores, limiting routine multi-color SMLM to two or three colors. Also, interpretation of hundreds of individual sites is challenging. Here we show how to bypass this limit using LocMoFit by reconstructing average density maps of multi-protein assemblies from pairs of dual-color data (Fig. 4a-d). In this strategy, we use one protein as a ‘reference structure’ that is always imaged together with a second ‘target’ protein, labeled in a different color. By fitting a model to the reference structure, we can determine the precise location and orientation of each site and thus register all sites from individual data sets, but also from data sets for different target proteins. Here we showcase this approach on the example of the NPC using endogenously tagged Nup96 as the reference structure (Fig. 4a-d). As target proteins we chose immunolabelled Elys, Nup133, Nup62, Nup153, and the NPC channel, stained by wheat germ agglutinin (WGA). From a fit of the NPC model (Fig. 2c) to the Nup96 localizations in all data sets and sites we could calculate average distributions of all target proteins (Fig. 4c) from single sites (Fig. 4a,b) and integrate all target proteins into a single coordinate system as an average protein density map (Fig. 4d). Note that this approach greatly increases the effective labeling efficiency of the target protein and can produce high-contrast averages even for very poor labeling (compare Fig. 4b to the average), and thus can reveal structural details not apparent in single images.

**Figure 4.**
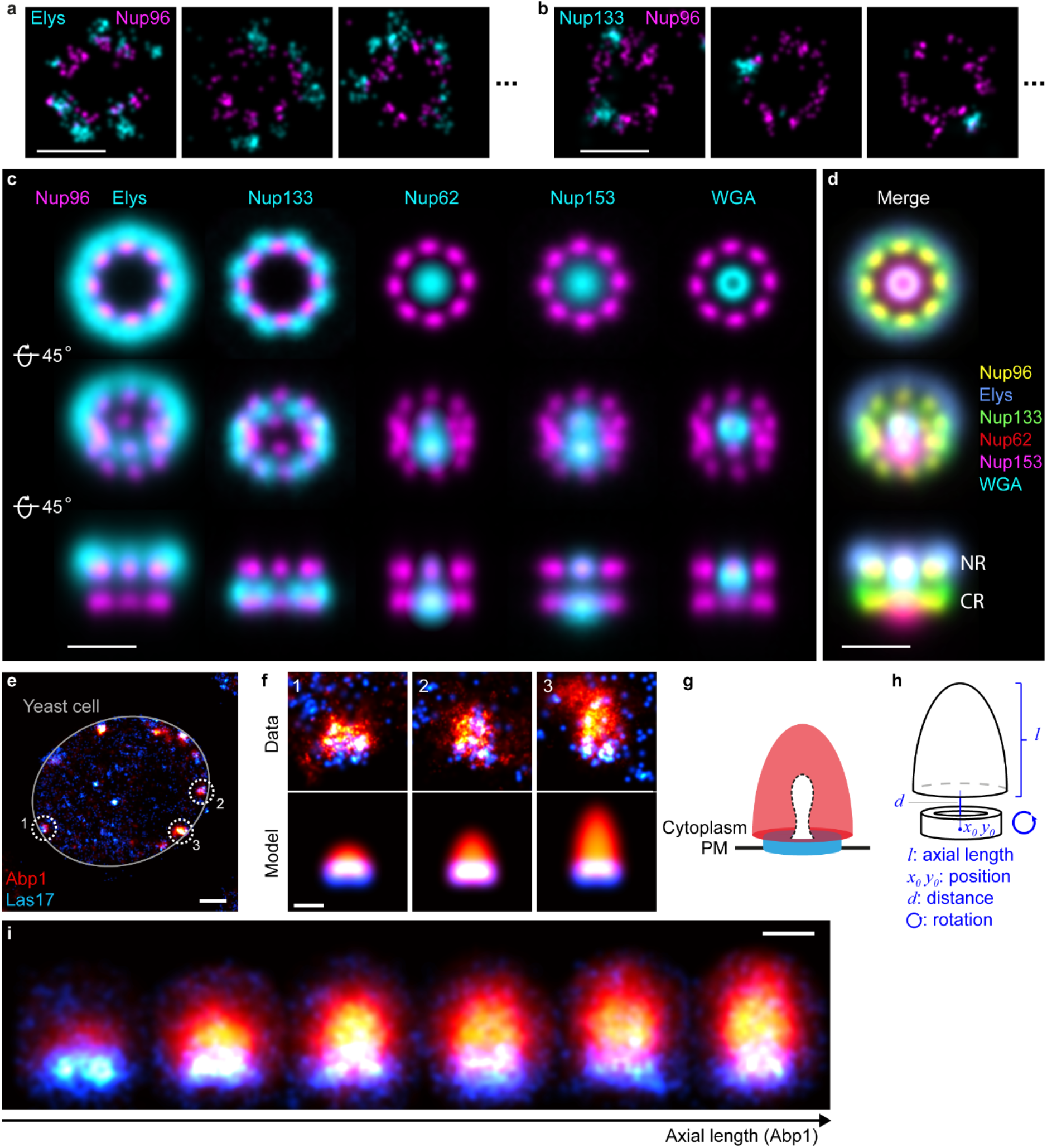
Static and dynamic protein density maps. **a-d, Average density map of the nuclear pore complex. a-b,** representative images of individual sites showing Nup96-SNAP-AF647 and immunolabeled Elys-CF680 (**a**) or Nup133-CF680 (**b**). **c,** A model fit to the reference protein Nup96 allows registering all sites of one data set and integrating different dual-color data sets into one common coordinate system (**d**). NR and CR denotes nucleoplasmic and cytoplasmic rings, respectively. See Supplementary Movie 1. Sample size: Elys: *n_s_* = 704; Nup133: *n_s_* = 704; Nup62: *n_s_* = 605; Nup153: *n_s_* = 419; WGA: *n_s_* = 488. *n_c_* = 1 for all. **e-i, Dual-color dynamic reconstruction of endocytosis in yeast. e,** Overview image of a single yeast cell showing Abp1-mMaple and Las17-SNAP-AF647. **f**, Individual endocytic sites are fitted with a dual-color model (**h**) that reflects the expected distribution (**g**) of Abp1 and Las17^6^: We model Abp1 as a semi-ellipsoid and Las17 as a thick ring and project these geometries in 2D. The fitted axial length of Abp1 is used as a proxy for pseudotime to sort individual endocytic sites according to their progression along the endocytic time line. The fitted position and orientation are then used to average all sites in each time bin (**i**). Bin size: 21 sites. Sample size: *n_s_* = 130, *n_c_* = 51. A running average can be found in Supplementary Movie 2. Scale bars: 100 nm (**a-d,f,i**), 500 μm (**e**).

LocMoFit can also perform a dynamic reconstruction of multi-protein assemblies from static super-resolution snapshots. The idea is to use LocMoFit to extract features of the structure that can be used for pseudo-temporal sorting and to then average individual structures in each time bin. We illustrate this approach on the example of the machinery that drives clathrin-mediated endocytosis in yeast, which is known to show highly regular dynamics and composition^32^. From prior super-resolution^6^ and electron-microscopy^33^ studies we know that the actin nucleation promoting factor Las17 forms a ring at the plasma membrane and that the actin binding protein Abp1 decorates the dome-shaped actin network that elongates during endocytosis^32^. By fitting a model that reflects this geometry to dual-color 2D data (Fig. 4e-h), obtained by focusing on the midplane of yeast cells, we use the length of the Abp1 structures to sort all sites according to their progression along the endocytic time line. We then distribute the structures in individual time bins and use the fitted position and orientation for averaging (Fig. 4i, Supplementary Movie 2) to result in dynamic protein localization maps.

### Model-free averaging

Model-free averaging or particle fusion is an approach that fuses particles, or sites, that share the same underlying structure together to form an average that approximates the underlying structure. Particles can be aligned based on templates/models, which however can lead to a so-called ‘template bias’: The final average can be biased towards the template, and even wrongfully reveal structures present in the template that are not present in the particles. Therefore, we would caution against interpreting the distribution of the reference Nup96 in Fig. 4. As the alignment of the other proteins was solely calculated based on this reference structure, they do not display template bias. Model-free particle fusion on the other hand allows calculation of unbiased average protein distribution maps and has been implemented based on alignment of particles using pair-wise cross-correlations^18,19,23^.

In LocMoFit we can use individual particles as models for other particles to determine their relative position and orientation and use those in an iterative workflow for model-free particle fusion (Fig. 5). Here, we illustrate this based on Nup96 in the NPC. As the log-likelihood is a measure for the similarity, we can efficiently construct the initial template: From an all-against-all pairwise registration of a subset of particles we can identify the particle that shows the highest degree of similarity to all other particles as a seed (Fig. 5b). We then cumulatively fuse the other sites in the order of their total similarity (Fig. 5c). This initial template is then used to register the remaining particles in the data set. The resulting average can then be used for the next round of registration (Fig. 5d). This step is iterated until convergence of the optimization (Fig. 5f).

**Figure 5.**
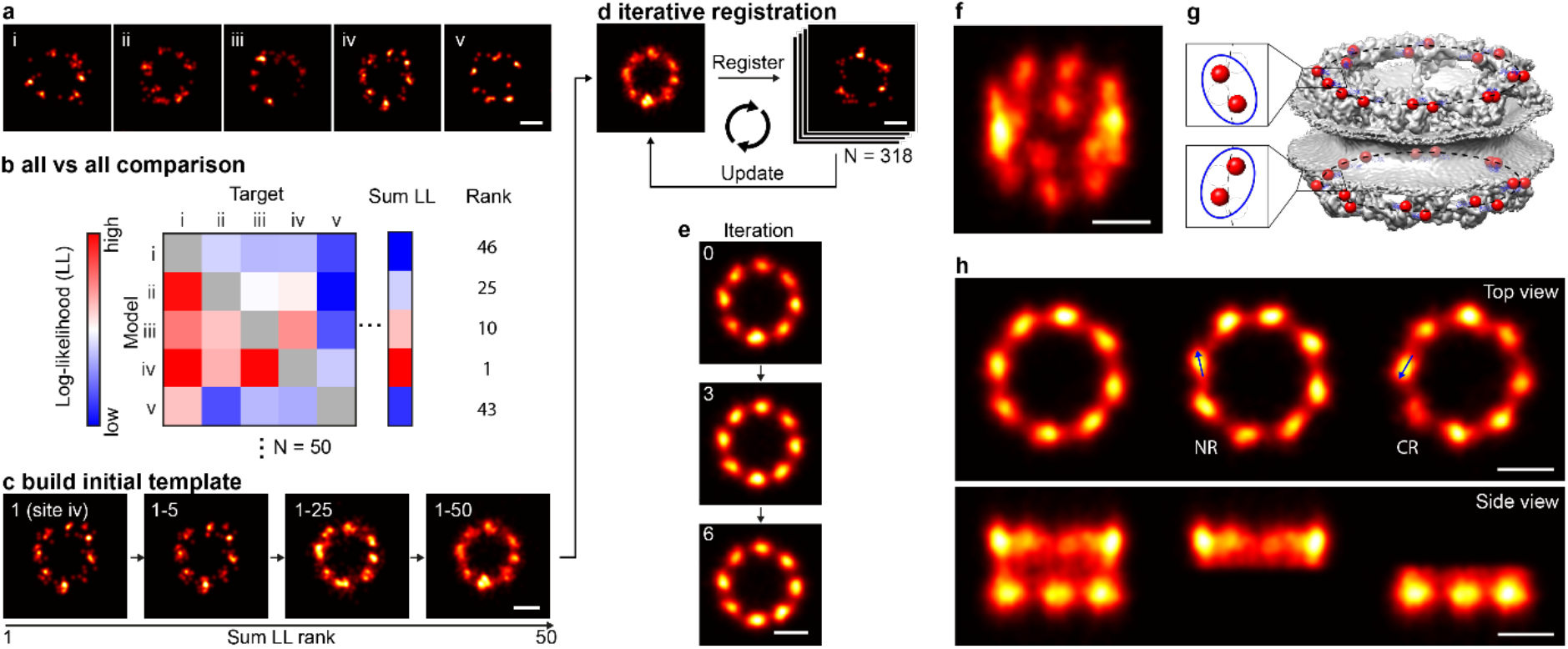
Model-free particle averaging. **a-e, Workflow. a**, Example single NPC particles. We assumed that all the sites are samples of the same underlying distribution. **b**, We first find a site (in the example the site iv) that best describes all other sites based on the rank on sum log-likelihood (LL) of the all-to-all matrix, where the 50 subset sites were fitted to each other. **c**, The initial template is built based on sequential registration in the order of the sum LL rank. **d**, The final fused particle is used to register all sites in the 318-site data set. This procedure yields an updated fused particle, which is used to register the data set again. This process is iterated until convergence. **f-h**, The final average calculated from 318 particles without any assumption on the underlying geometry or symmetry in a tilted view (**f**), and for comparison the EM density of the NPC with C-termini of Nup96 indicated in red. (**g**), Top and side view, where the nucleoplasmic and cytoplasmic rings are shown together (left panel), or separately (middle, right panels) (**h**). See Supplementary Movie 3. The two proteins per ring per symmetric unit give rise to tilted elongated average protein distributions in the averages (arrows in **h**). Scale bars: 50 nm.

The resulting 3D average of Nup96 clearly resolves the two rings in the NPC and their 8-fold symmetry (Fig. 5g). In addition, it reveals subtle structural details such as the elongated, tilted shape of the corners (Fig. 5h) that directly reflects that in each ring each symmetric unit is occupied by two Nup96 copies with slightly different radii (Fig. 5g).

In summary, LocMoFit allows for bias-free high-quality 3D averaging without any assumptions on the underlying geometry and symmetry.

## Discussion

In this study, we present LocMoFit, a powerful and general framework for extracting quantitative descriptors of cellular structures by fitting an arbitrary, parameterized model to SMLM data. As the fitting is performed on individual structures without averaging, this will help investigating biological heterogeneities among structures that are currently difficult to quantify with electron microscopy, where typically many identical structures need to be averaged to reach sufficient signal-to-noise ratios.

LocMoFit relies on choosing a model that can represent the data. A wrong model will still result in the most likely parameters that explain the experiment, but these parameters then might become difficult to interpret or meaningless. So how can we construct a model for a biological structure? Usually, a simple geometry or symmetry can be guessed by visual inspection of the data, from prior knowledge based on other techniques, or by model-free particle averaging, as also implemented in LocMoFit. It is then key to define the parameters in a way that the model is as general as possible and can describe a large class of experimental structures. For instance, all the models used in this study are not rigid templates, but can change their size and shape during optimization. In case of competing models, the more likely model can be chosen based on its lower AIC_C_ (Fig. 3), taking the sample size and the number of free parameters into account^22^.

Of equal importance to selecting the right model is the quality of the data, which must contain sufficient information to unambiguously define the multiple model parameters. In case of low labeling densities, large localization errors or structures with few features, simple models with few free parameters have a lower risk for overfitting than complex models. Even a well-chosen model might not converge to the global optimum. In these cases, choosing proper initial parameters in a first fitting step with a simpler model or even manually can provide a good solution, as choosing an optimizer in LocMoFit that performs a parameter search over defined intervals instead of gradient descent.

LocMoFit is equipped with tools for facilitating the reliability of data analysis. One of the tools is visualization, which allows users to inspect, as we always recommend, the results of the fit efficiently. Furthermore, the reliability is hard to examine without knowing the ground truth of the data^34,35^. In line with this, LocMoFit provides a simulation engine that generates SMLM coordinate data with a given model and known parameters and allows investigating the precision of the parameter estimates, the suitability of a model to fit data of a specified quality and the impact of initial parameters on convergence.

Integrating large datasets into an average representation in the form of protein localization maps can be a useful and complementary approach to a statistical analysis of parameters extracted from individual sites. LocMoFit can calculate such average representations by determining the precise position and orientation of a reference structure and use this to align target proteins, imaged in a second channel. By additionally evaluating a parameter that changes monotonously over time, LocMoFit can extend this approach to dynamic, time-resolved localization maps. However, whenever templates or models are used for registration, the averages might be biased towards the model^36^. Thus, care should be taken when interpreting the averages of the reference structures. Target structures that were not considered during the alignment, on the other hand, should be free of such model bias.

This model bias can be avoided largely when using template-free particle averaging (Fig. 5). This approach is commonly used in electron microscopy and has been introduced to SMLM data recently^23,18,19^, with or without an adaption. The adaption was implemented because of the different data types between SMLM (sparse and coordinate-based) and EM (dense and intensity-based)^18^. LocMoFit too performs particle fusion directly on the coordinates and takes into account the localization precisions. Our new approach of selecting the most representative particles to construct the initial template minimizes the bias of seed selection, while avoiding a computationally expensive all-to-all registration.

When we construct average protein density maps or template-free particle averages, we need to assume that all particles are based on the same underlying structure. However, biological variability is expected in most experiments. In this case, the resulting distributions are averages over the different conformations and can be biased towards a sub-population. In the future, a combination of averaging with classification^24^, as we demonstrated when we reconstructed the dynamic protein maps of endocytosis (Fig. 4i), could extend particle averaging to heterogenous and dynamic cellular structures.

In the current framework, more than one fluorophore per target molecule and repeated activation of a single fluorophore are not considered during fitting, but we investigated their impact with simulations. A future extension to a probabilistic model of fluorophore blinking and non-stoichiometric labeling, possibly using a Bayesian framework^37^, would further improve robustness and accuracy.

Currently, a fit to a single site takes seconds to minutes (5-10 seconds for an NPC and CME site, 10-20 minutes for a micrometer-long microtubule, depending on the complexity of a model and the number of localizations of a site), allowing even large data sets with hundreds of sites to be analyzed in overnight runs on a standard CPU (e.g., Intel core i5-4460). In the future, deploying LocMoFit on clusters or GPUs could further improve performance.

Published as open-source, LocMoFit is readily useable as part of the SMLM software platform SMAP^21^, allowing users to easily fit their own data with any of the numerous pre-defined models using a graphical user interface. To this end, we provide detailed documentation, tutorials, and example files. Alternatively, LocMoFit can be run independently of SMAP and provides an API for integration in own software. All models used in this study are ready-to-use and available to the public domain and can be combined into complex composite models. New models can be created with basic programming expertise. We encourage users to deposit their own models to our Git repository to facilitate knowledge sharing.

LocMoFit will enable many researchers to greatly increase the information that can be extracted from their data and to develop new and complex data analysis workflows that drive biological discovery.

## Methods

### LocMoFit framework

#### Model fitting in LocMoFit

LocMoFit fits a parameterized geometric model to a set of localizations from the same site through Maximum Likelihood Estimation (MLE). LocMoFit requires two inputs: 1) a parameterized geometric model *f*(*p*) that describes the distribution of the fluorophores in the structure with the set of parameters *p* of the model, and 2) a set of *K* localizations 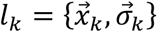. 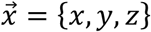 are the coordinates of a detected emitter and 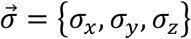 the associated uncertainties. 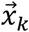 and 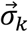 are typically obtained by fitting an experimental or Gaussian point spread function model to the raw camera frames using maximum likelihood estimation^38,39^.

To take into account localization uncertainties, we do not use *f*(*p*) directly for fitting, but instead a probability density function (PDF) 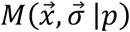, which is derived from *f*(*p*) as described in the next section. 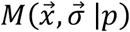 describes the probability of finding a single random localization *l* at the coordinate 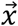 given an uncertainty 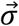 and model parameters *p*.

If we measure a set *l_k_* of *K* localizations and assume that they are random and independent variables of the PDF 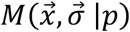, the likelihood to obtain precisely these localizations *l_k_* is simply the product of individual probabilities as shown in equation (1).

To find the set of parameters 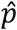 that, together with *M*(*p*) and therefore *f*(*p*), best describes *l_k_*, we maximize this likelihood using an optimization algorithm (see section **Optimization procedure**) as shown in equation (2).

#### Calculation of the probability density function

Here we discuss how to calculate the PDF 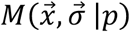 from the geometric model *f*(*p*). *f*(*p*) can be defined as either a *fluorophore density map, discrete fluorophore coordinates* or a *continuous fluorophore distribution*.

In the first scenario when defined as a *fluorophore density map*, the geometric model 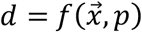 directly outputs the density *d* of the fluorophore at the position 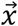. Here 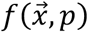 is not necessarily normalized. If the localization uncertainty has been included in *f*(*p*), its PDF 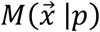 can be derived by simple normalization:

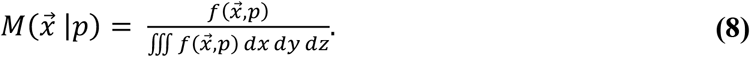

Otherwise, the uncertainty can be introduced by convolving 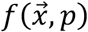 with a Gaussian function with a standard deviation defined by the average localization uncertainty 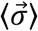 as in equation (3). In practice, the model *f*(*p*) can be supplied as an image for a 2D fit, an image stack for a 3D fit, or directly as a function.

In the second scenario when defined as *discrete fluorophore coordinates*, the geometric model *f*(*p*) specifies the expected coordinates *v_j_* of the fluorophores so that *v_j_* = *f*(*p*). To derive the PDF for this case, let us consider a simple one-dimensional example, in which a fluorophore at position *v* with a localization precision *σ* is repeatedly localized, resulting in measured coordinates *x_k_*. These measured coordinates then scatter around the true position with a standard deviation of *σ*, following a Gaussian distribution. Thus, the probability that the measured coordinate *x* is caused by the fluorophore at position *v* is^18,23,19^:

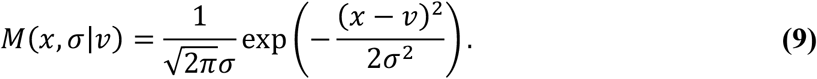

If we have *J* model fluorophore positions 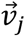, the probability that they describe a single measured localization 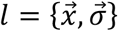 is given by the sum of the individual probabilities (now for the 2D or 3D case) as in equation (4). The likelihood function *L*(*p*) is then calculated according to equation (1) by multiplying the probabilities of all measured localizations. Compared to the first case (equation (3)), in which only the average localization precision is used to blur the model, here all localization precisions 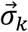 contribute individually to the PDF so that more precise localizations have higher impact. As the stochastic nature of single-molecule imaging leads to a wide distribution of the localization precisions, this increases the accuracy by properly weighting the single localizations in the PDF.

In the third scenario when defined as *continuous fluorophore distributions*, the geometric model 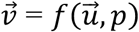 describes a parametric line (e.g. a filament or a ring) or surface (e.g. a spherical shell), in which fluorophores are distributed with constant density. The vector variable 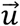 has only one element when describing a parametric line and two when describing a surface. In practice, LocMoFit works with a discrete form *f_d_*(*p*) of the geometric function 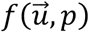. To discretize 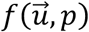, either LocMoFit renders *J* fluorophores 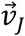 on the line or surface based on *J* vectors 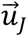 across the range defined by the user and assigns every point a weight *q_j_* inversely scaled to local density, having 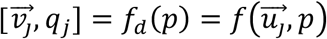, or the user defines *J* fluorophores 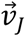 evenly distributed on the line or surface defined by 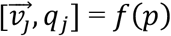, with *q_j_* = 1. In either case, the maximum spacing *δ* between adjacent points defined in 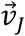 is required to be smaller than the minimal localization precision of the *K* localizations to retain continuity: 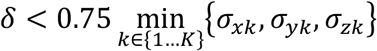. To improve the computational speed by reducing the size of the model, LocMoFit also allows the user to define a minimal localization precision *σ_min_* so that any *σ_xk_, σ_yk_*, and *σ_zk_* smaller than *σ_min_* are set to *σ_min_*. This setting increases the required spacing *δ* and reduces the required sampling rate (associated with *J*) and therefore the size of the model. With discrete positions of fluorophores 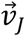, the convolution can be seen as placing Gaussian functions centered at all the positions in 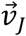. By having 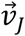, we can utilize equation (4) to construct the PDF with the introduction of *q_j_*:

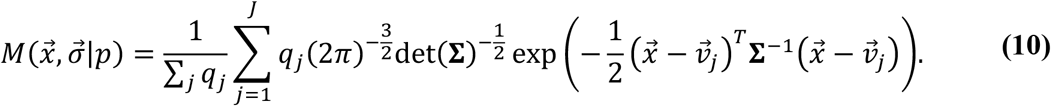

Equation (4) is then a special form of equation (10) with *q_j_* = 1.

In this study, we refer to a discrete model when it is constructed based on either equation (4) or equation (10), and to a continuous model when it is constructed based on either equation (3) or equation (8).

#### Optimization procedure

To find the set of parameters 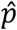 that maximizes *L*(*p*), the user can select either an evolutionary algorithm that searches parameters more globally, a simplex-based derivative-free searching, or a gradient-descent optimizer. Before optimization, the user can define which parameters to fit and which to set to a constant value, and their initial guesses and boundaries. The initial guesses can be either pre-defined values or values derived from user-defined rules.

For fitting, we classify the parameters *p* into intrinsic parameters *p^i^* that directly determine the shape of the model and extrinsic parameters 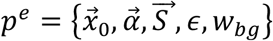 that describe the position of the model 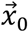, the orientation, described by the rotation angles 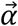 about the three axes, and optionally a global scaling factor 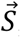, an uncertainty *ϵ* additional to the localization precision, and the weight *w_bg_* of a constant background PDF *M_bg_* to accommodate the localizations that cannot be described by the geometric PDF (see next section). Here the rotation angles 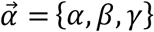 about the *x, y*, and *z* axes, respectively, define the rotation matrix:

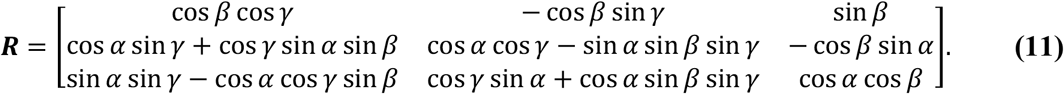

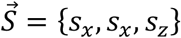 contains the scaling factors of the three spatial axes, defining the scaling matrix 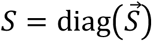. For a model in the continuous form, we use the extrinsic parameters *p^e^* to reversely transform the localizations, which is computationally more efficient than to transform the model. Thus, during the optimization, we first transform the localization coordinates as

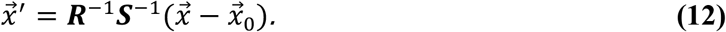

For a discrete model we translate and rotate the model instead to avoid computationally costly rotation of the anisotropic multidimensional Gaussian (equation (4)), particularly in 3D. In this case, the fluorophore positions of the model 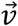 are transformed during optimization as:

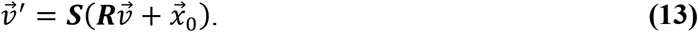

As a result of maximizing the likelihood with respect to *p^i^* and *p^e^*, we obtain the parameter estimates 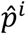 and 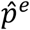 along with their 95% confidence intervals.

#### Background localizations and additional uncertainties

In real experiments, unspecific background fluorophores, localizations from neighboring structures or large localization errors lead to localizations that are not described by the model. This mismatch can introduce a bias in the parameter estimates. We accommodate these so-called ‘background’ localizations with an evenly distributed (constant) PDF *M_bg_*:

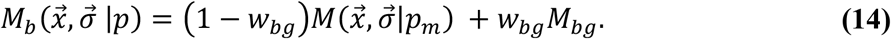

The set of parameters *p_m_* contains all elements of *p* except for the background weight *w_bg_*. *M_bg_ = *d*^−D^* with *d* the length of a site and *D* is the dimension, so that the summed probability of *M_bg_* over the site is one. *w_bg_* is the background weight that represents the fraction of localizations that are considered background. The total number *K_bg_* and density *ρ_bg_* of background localizations can be obtained as *K_bg_* = *K* · *w_b_* and *ρ_bg_* = *K_bg_*/*d*^2^. The difference of the total number of localizations *K* and *K_bg_* is then the total number of localizations described by the model *K_m_* = *K* – *K_bg_*.

The localization precision *σ* often underestimates the true spread of localizations in real experiments. The reason can be instabilities like thermal drifts or vibrations during the experiment, the size of the label that displaces the fluorophore from the target structure (linkage error) or biological variability that leads to a spread of the fluorophores that is not described in the model. These additional uncertainties, quantified by the parameter *ε*, lead to an additional blurring (equation (3)) with 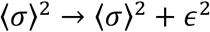. In equation (4) we take *ε* into account with a modified covariance matrix:

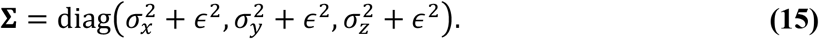

*ϵ* can be specified by the user or used as an additional free fitting parameter.

#### Composite models

LocMoFit allows the user to combine several simple models into a single one by adding up and re-normalizing the PDFs of each model (see equation (5)).

The sum of weights is one: ∑_*m*_ *w_m_* + *w_bg_* = 1. *w_m_* represents the proportion of the localizations that can be described by the component PDF *M_m_*. With the weights we can estimate the number of localizations *K_m_* coming from a specific component model *M_m_* by *K_m_*=*K* · *w_m_*.

Note that here we define the extrinsic parameters 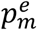 (except for the model weight *w_m_*) of the *m^th^* component model (*m* > 1) with respect to the first component model, with a value zero indicating the same transformation as the first component model. That is, the rigid transformation of the first component model (according to 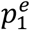) is first applied to all component models, followed by the rigid transformation of the *m^th^* component model (according to 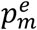) applied to only the *m^th^* component model.

The user can select which parameters are fixed among the models and which are fitted independently. This greatly facilitates constructing complex models.

When fitting multi-color SMLM data, each localization is not only described by its coordinate and localization precision, but also its color *c*. In this case, we can define a separate model for each color channel and fit all models simultaneously, as shown in equation (6).

The weight for each color channel *w_c_* is introduced to minimize the effects of different numbers of localizations between different colors and can be assigned as *w_c_* = ∑_*c*_ *K_c_*/*K_c_*, where *K_c_* is the number of localizations with the color *c*. *w_c_* is used as an exponent to normalize the different multiplications, which scales to the number of localizations, in Equation (1). When the effects of different numbers of localizations are preferred, weighting can be switched off by setting *w_c_* = 1. Note that each single-color PDF 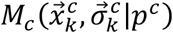 (with the background PDF 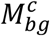, as described by equation (5) for an individual model or equation (6) for a composite model) is evaluated only with the localizations of the corresponding color.

#### Chaining fitting steps for improved convergence

For complex models with many fitting parameters, optimizers are limited in scanning the entire parameter space to find a global optimum and might get stuck in a local maximum of the likelihood. Thus, LocMoFit allows the user to chain several fitting steps with different models and use the results of the previous step as the initial parameter for the next one. Note that the first step can employ user-defined rules/functions to provide initial parameter estimates. Then, the user can use a less complex model with strong blur (equation (15)), using a global optimizer before fine-tuning the fit with a simplex or gradient-descend optimizer on the precise model. In this way, LocMoFit efficiently finds the global maximum of the cost function *L*(*p*).

#### The relation between likelihood and cross-correlation

The likelihood *L*(*p*) can be seen as a metric that describes the similarity between model *f*(*p*) and data *l_k_* from the probabilistic aspect. By changing the multiplication in equation (1) to summation, we get another metric that is regularly used for pattern matching and represents the cross-correlation between model and data:

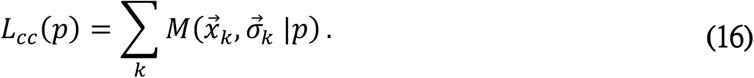

When using a model *f*(*p*) in the discrete form, by plugging its PDF (as in equation (4)) into equation (16), we get a similar form as the correlation between two sets of points derived in Schnitzbauer et al.^23^, with the exception that we do not assign uncertainties to fluorophore coordinates in the model. Also, it is closely related to the Bhattacharya cost function and derivations that were previously used for particle fusion^18,19^ and detecting structural heterogeneity^24^ in SMLM. Therefore, the cross-correlation *L_cc_*(*p*) can also be used as the objective function in LocMoFit.

### Data analysis

#### Model fitting

Model fitting requires segmented sites (see the section **Segmentation of sites**).

##### Nup96

We used three models to describe Nup96 in different fitting steps. The first model, *NPC*_*m*1_, is a composite model of two identical rings, with a radius *r* of 53.7 nm, shifted along their common axis. The extrinsic parameters of the upper ring were fixed to those of the lower ring, except for the *z* position. This model was implemented as a fluorophore density map. The second model, *NPC*_*m*2_, is a dual-ring model that has two identical parallel rings, parameterized by intrinsic parameters ring radius *r* and ring separation *s*. This model was implemented as a discretized continuous fluorophore distribution. The third model, *NPC*_*m*3_, was built using the *NPC*_*m*2_ as a backbone, having the continuous rings replaced by fluorophore positions (Fig. 2c). Two of the fluorophores form a unit, which is evenly placed eight times on one ring rotationally and yield 32 positions in total. Since the rings are not continuous anymore, the twist *θ* between the two rings is also an intrinsic parameter in addition to the two parameters inherited from the second model.

For single-color NPC data (Fig. 2a-f), we chained these three fitting steps: 1) fitting with *NPC*_*m*1_ to roughly measure the orientations, positions, and ring separations of the NPCs, 2) fitting with *NPC*_*m*2_ to refine the previously measured parameters and to measure radii, and 3) fitting with the *NPC*_*m*3_ to measure the ring twist, with the extra uncertainty *ϵ* a free parameter to allow exploring parameter space more during optimization. The initial parameters of a later step were inherited from the final parameters of the previous step. All parameter settings are summarized in Supplementary Table 3.

For dual-color NPC data (Fig. 4a-d), Nup96 was fitted in two chained steps: 1) fitting with *NPC*_*m*1_ as for the single-color data, and 2) fitting with *NPC*_*m*3_, having intrinsic parameters fixed to the mean parameter values that were extracted from the single-color data (Fig. 2d-f). All parameter settings are summarized in Supplementary Table 5.

For the model selection, the fitting steps were the same as for single-color NPC data except that different rotational symmetries were used as specified.

##### Microtubules

We used two models to describe microtubules. The first model, *MT*_*m*1_, describes a cubic spline in 3D. In this model, the spline is defined as piece-wise third-order polynomials that traverse through a set of odd number *N* of equidistant control points, in the order *q* = 1 to N. The middle point (*q* = *q*_0_ = (*N* + 1)/2) is defend as the reference position 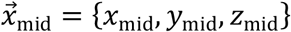. Starting from the middle point, the rest of control points are defined in two directions, one from *q* = *q*_0_ – 1 to 1 and the other from *q* = *q*_0_ + 1 to *N*. Following these orders, the position of one control point (*q* = *q_n_*) is defined by its distance *d* from the previous control point and the azimuth *θ_q_* and elevation angle *φ_q_*, defined relative to the previous control point. The second model, *MT*_*m*2_, uses the first model as a backbone, rendering rings, centered at equidistant points on the backbone spline, perpendicular to the backbone (Fig. 2i). Thus, the radius *r* of the rings is an intrinsic parameter in addition to the ones inherited from the first model. Both models were implemented as discretized continuous fluorophore distributions. In this study, we used the number of control points *N* = 5 and the distance between points *d* = 250 nm.

Microtubules (Fig. 2g-j) were fitted with two chained steps: 1) fitting with *MT*_*m*1_, having a large extra uncertainty *ϵ* to estimate the center line of microtubule segments, and 2) fitting with *MT*_*m*2_ to refine the path of the microtubules and to measure the radius. The initial parameters of the second step were inherited from the final parameters of the first step. All parameter settings are summarized in Supplementary Table 4.

##### Endocytic structures

For fitting endocytic sites, we used a composite model formed by a two element model: projections of a 3D hemispheroid and a thick ring onto the 2D imaging plane (Fig. 4g). This model was implemented as a discretized continuous fluorophore distribution. In the imaging plane, the base of the hemispherical projection is limited to below the thick-ring projection. The hemispherical projection is parameterized by the half long and half short axes *a* and *b* of a hemispheroid. The thick-ring projection is parameterized by the thickness *t* and the inner/outer radii *r* and *q* of the ring. This model was fitted to the yeast endocytic sites in the dual-color dataset (Fig. 4e-i). The hemispheroid was only fitted to Abp1 localizations and the thick ring to Las17 localizations, respectively. All parameters are summarized in Supplementary Table 6.

#### Simulation

We performed realistic simulations based on a two-state (bright and dark) fluorophore model plus bleaching^40^. (1) First, we defined model parameters, which can be fixed numbers or uniformly distributed random variables within specified boundaries. (2) With the defined model parameters, we generated protein positions for each simulated site by taking all the *N* positions (e.g, 32 positions for the eight-fold symmetric model of the NPC) of proteins defined in a point model or randomly drawn *N* samples from a specified PDF with no uncertainty. (3) With a probability *p*_label_, a fluorescence label was created at a protein position. (4) Linkage displacements in *x, y* and *z* were added to a label and were determined as normally distributed random variables with a variance corresponding to the linkage error *ϵ*. (5) Each fluorophore appeared at a random time and lived for a time *t_l_*, determined as a random variable from an exponential distribution. (6) A label had a probability *p*_react_ to be reactivated and then appeared at a random later time point, otherwise it was bleached. (7) When it was on, a fluorophore had a constant brightness. Thus, the brightness in each frame was proportional to the fraction of the time the fluorophore was on in each frame. (8) The emitted photons in each frame were determined as a random Poisson variable with a mean corresponding to the average brightness in the frame. (9) For each frame, we calculated the CRLB (Cramér-Rao lower bound) in *x, y* and *z* from the number of photons and the background photons^41^. (10) This error was added to the true *x, y* and *z* positions of the fluorophores as normally distributed random values with a variance corresponding to the respective calculated CRLB. Simulation parameters are summarized in Supplementary Table 1.

The simulated localizations were processed with the same data analysis pipeline as the real data.

#### Reference-based averaging of multi-color data

To create the average density map of the nuclear pore complex, in each site only Nup96 localizations were fitted as described in the section **Model fitting in LocMoFit, *Nup96***. Each site was transformed to the orientation and position of the model so that all the sites were in the same coordinate system. The transformed localizations of all sites then formed the averages.

For the dynamic reconstruction of CME in yeast, all sites were sorted by the fitted length of the hemi-ellipsoid describing Abp1 localizations. The orientation of each site was aligned to the direction of the membrane invagination and the estimated position of the Las17 ring model defined the origin. Each time bin was then created from the localizations of 21 aligned sites. The movie of the dynamic reconstruction (Supplementary Movie 3) was generated by running averaging through the aligned sites over the pseudotime: each frame comprised 15 sites and the step size was one site.

A technical limitation of this example is the use of indirect immunolabeling. Here, varying epitope accessibility and non-random orientation of the antibodies can result in systematic differences between protein density maps and true distributions of the proteins, which in principle can be overcome with improved labeling schemes.

#### Model selection

In LocMoFit, we provided the corrected Akaike information criterion^22^ (*AIC_C_*) as the metric for performing the model selection. In general, a model with more free parameters tends to fit better. Therefore, instead of using the maximum likelihood 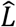 as the metric, 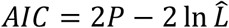 was suggested for penalizing the number of free parameters *P*^22^. In practice, we would like to choose a model with fewer parameter and yet a larger maximum likelihood. Therefore, the smallest AIC indicate the best model when fitting the same data. To avoid overfitting caused by small sample size, the corrected *AIC_C_* includes an additional penalty: *AIC_C_* = *AIC* + (2*P*^2^ + 2*P*)/(*K* – *P* – 1), where *K* is sample size^22^. When *K* → ∞, the additional penalty term approaches zero so that *AIC_C_* converges to *AIC*. In LocMoFit, the sample size *K* is the number of localizations. For visual comparison of the *AIC_C_* we normalize by the number of localizations *K*.

#### Model-free averaging

For model-free averaging of Nup96 particles, we first estimated the approximate orientations and positions of each particle by fitting with a continuous dual-ring model. Next, we generated an initial model from a subset of transformed particles *P*_1_… *P*_50_. To this end, we defined the localization coordinates of each particle as the fluorophore positions of a point model. Positions and orientations of all model particles were corrected according to their estimates. Then, we fitted each model to all other particles in the 50-particle subset. Based on LL values acquired by all-against-all pairwise fitting, we then built a similarity matrix *M*. We then cumulatively fused the particles in the order *R* of their total similarity: each particle *P*_[*R*=*i*]_ was registered to the fused particle *T*_[*R*=*1*−1]_ starting with the highest-ranked particle *T*_1_ = *P*_[*R*=1]_. This initial model *T* that was used to register the remaining particles *P*_51_… *P*_200_ in the 200-particle data set. The resulting average *T* was then used as the new initial template for the next round of registration. This step was iterated until convergence and yielded the final average 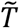. See **Algorithm 1** for the pseudocode.

##### Algorithm 1. Model-free averaging

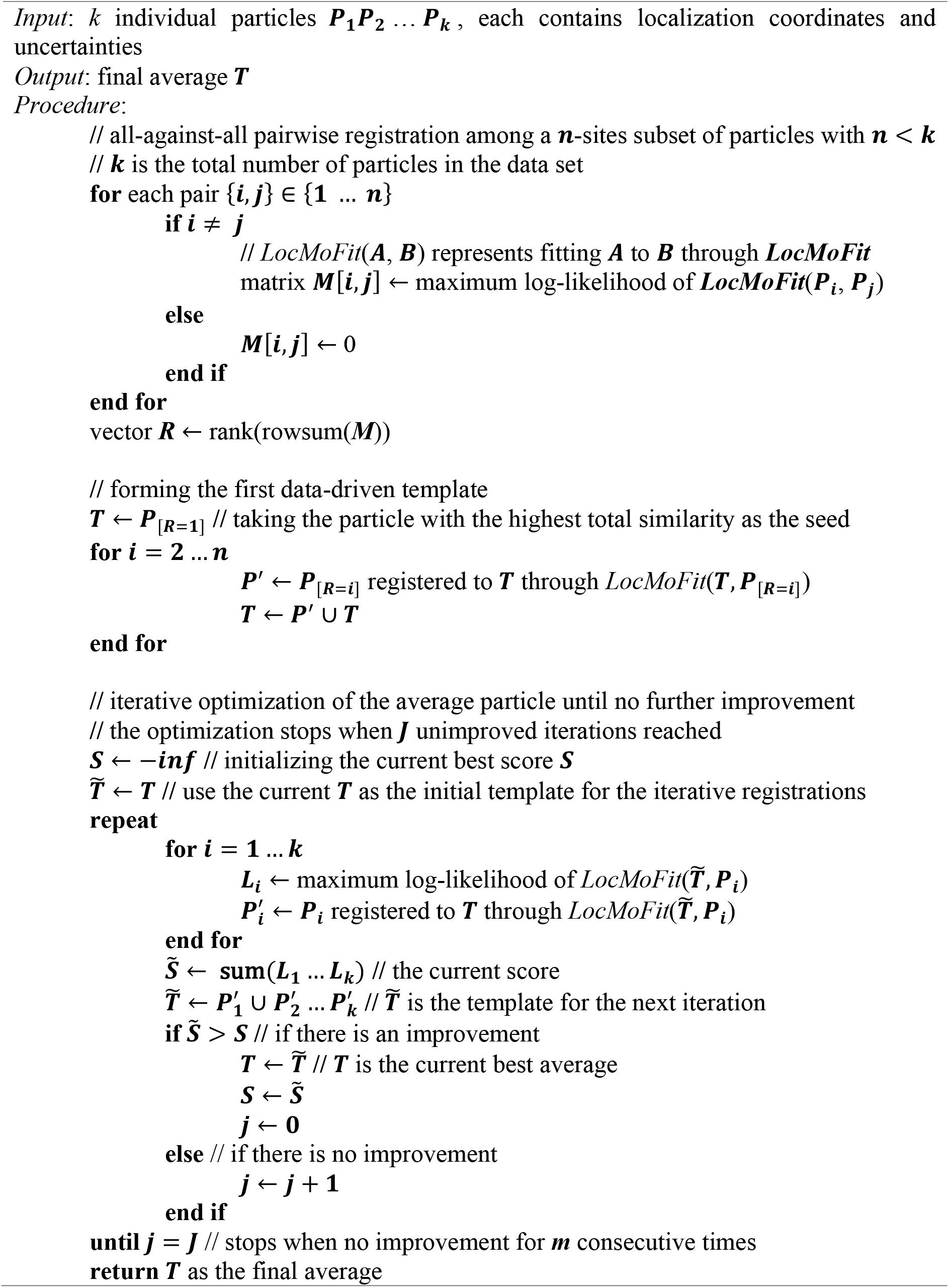

### Sample preparation

#### Preparation of coverslips

24 mm round glass coverslips were cleaned overnight in methanol/hydrochloric acid (50/50) while stirring. They were then rinsed repeatedly with ddH2O until the pH of the washing solution remained neutral. They were then placed overnight into a laminar flow cell culture hood to dry them before finalizing the cleaning of coverslips by ultraviolet irradiation for 30 min.

For yeast samples, the coverslips were subsequently plasma cleaned for 5-10 min. A drop of 20 μL concanavalin A (ConA) solution (4 mg/mL in PBS) was added to each coverslip, spread out with a pipette tip, and let incubate for 30 min in a humidified atmosphere. Then, the remaining liquid was removed and the coverslips were dried overnight at 37°C. Prior to use, the remaining salts were washed off with ddH_2_O.

#### Sample seeding

Cells were seeded on clean glass coverslips 2 days before fixation to reach a confluency of about 50–70% on the day of fixation. They were grown in growth medium (DMEM; catalog no. 11880-02, Gibco) containing 1× MEM NEAA (catalog no. 11140-035, Gibco), 1× GlutaMAX (catalog no. 35050-038, Gibco) and 10% [v/v] fetal bovine serum (catalog no. 10270-106, Gibco) for approximately 2 days at 37 °C and 5% CO2. Before further processing, the growth medium was aspirated and samples were rinsed twice with PBS to remove dead cells and debris. Unless otherwise stated, all experimental replicates were performed on cells of a different passage with separated sample preparation.

#### Imaging buffers

Yeast samples were mounted in D_2_O blinking buffer (50 mM Tris-HCl pH 8, 10 mM NaCl, 100 U/mL glucose oxidase, 0.004% [w/v] catalase, 10% [w/v] D-glucose, 20 mM cysteamine, in 90% D_2_O).

NPC samples were imaged in 50 mM Tris-HCl pH 8, 10 mM NaCl, 100 U/mL glucose oxidase, 0.004% [w/v] catalase, 10% [w/v] D-glucose, 35 mM cysteamine.

#### Preparation of NPC samples

For single-color imaging, coverslips containing Nup96-SNAP-tag cells (catalog no. 300444, CLS Cell Line Service) were rinsed twice with warm PBS. Prefixation was carried out in a 2.4% [w/v] formaldehyde (FA) in PBS solution for 40 s before the samples were permeabilized in 0.4% [v/v] Triton X-100 in PBS for 3 min. Complete fixation was carried out in 2.4% [w/v] FA in PBS for 30 min followed by 3 × 5 min washing steps in PBS after fixation. Subsequently, the sample was incubated for 30 min with Image-iT FX Signal Enhancer (catalog no. I36933, Thermo Fisher Scientific) before staining with SNAP dye buffer (1 μM BG-AF647 (catalog no. S9136S, New England Biolabs) and 1 μM dithiothreitol in 0.5% [w/v] bovine serum albumin (BSA) in PBS) for 2 h at room temperature. To remove unbound dye, coverslips were washed three times for 5 min in PBS. At this point, the sample was ready for single color superresolution imaging.

For simultaneous dual-color imaging with immunostaining, samples were further blocked with 5% [v/v] normal goat serum (NGS) (catalog no. PCN5000, lifeTech) in PBS for 1 h. Binding of primary antibody (Elys (catalog no. HPA031658, Atlas Antibodies, 1:50), Nup133 (catalog no. HPA059767, Atlas Antibodies, 1:150), Nup62 (catalog no. 610498, BD Biosciences, 1:150), Nup153 (catalog no. ab24700, Abcam, 1:60)) was achieved by incubation with the respective antibody diluted in 5% [v/v] NGS in PBS for 1 h. Coverslips were washed three times for 5 min with PBS to remove unbound antibody and subsequently stained with CF660C labeled anti-rabbit antibody (catalog no. 20183, Biotium) or anti-mouse antibody (catalog no. 20815, Biotium) diluted 1:150 in PBS containing 5% [v/v] NGS for 1 h. After three washes with PBS for 5 min each, the sample was postfixed for 30 min using 2.4% [w/v] FA in PBS, rinsed with PBS, quenched in 100 mM of NH4Cl for 5 min and rinsed three times for 5 min with PBS.

For simultaneous dual-color imaging with WGA staining, cells on a coverslip were fixed, permeabilized, and stained with SNAP dye as described above. The sample was then incubated for 10 min with 1:5,000 diluted WGA-CF680 (catalog no. 29029-1, Biotium) in 100 mM Tris pH 8.0, 40 mM NaCl, and rinsed three times with PBS.

Before imaging, samples were mounted into custom sample holders in appropriate imaging buffers (see section Imaging buffers). The holder was sealed with parafilm.

#### Strain and sample preparation for yeast

The yeast strain expressing Abp1 tagged with mMaple^42^ and Las17 tagged with SNAPftag^43^ was described previously (JRY0014; ^6^). Briefly, the two proteins were tagged at their C-termini at the endogenous loci^44^. The strain was verified by colony PCR and fluorescence microscopy. Prior to the day of imaging, yeast cells were inoculated from single colonies on plates into 10 mL YPAD in a glass flask, and grown overnight at 30°C with shaking. The next morning, the culture was diluted into 10 mL YPAD in a glass flask to an OD_600_ of 0.25, and grown for 3 more hours at 30°C, typically reaching an OD_600_ of 0.6-1.0.

For sample preparation, 2 mL of the culture were collected by centrifugation at 500 rcf. for 3 min, resuspended in 100-150 μL YPAD, and pipetted on a ConA-coated coverslip. During all following incubation steps the samples were protected from light. The cells were allowed to settle for 15 min in a humidified atmosphere. Next, the coverslip was directly transferred into the freshly prepared fixation solution (4% [w/v] FA, 2% [w/v] sucrose in PBS). After 15 min of fixation with gentle orbital shaking, the sample was quenched in 100 mM NH_4_Cl in PBS for 15 min. Quenching was repeated once more before the coverslips was washed once in PBS for 5 min. Next, cells were permeabilized for 30 min by addition of the permeabilization solution (0.25% [v/v] Triton X-100, 50% [v/v] ImageIT FX, in PBS). The coverslip was washed twice in PBS for 5 min and then transferred face down on a drop of 100 μL staining solution (1 μM SNAP Surface Alexa Fluor 647, 1% [w/v] BSA, 1 mM DTT, 0.25% [v/v] Triton X-100, in PBS) on parafilm. After staining for 90 min, the sample was washed 3 times in PBS for 5 min each.

### Microscopy

#### Microscope setup and imaging

All SMLM data in mammalian cells were acquired on a custom built widefield setup described previously^6,45^. Briefly, the free output of a commercial laser box (LightHub, Omicron-Laserage Laserprodukte) equipped with Luxx 405, 488 and 638 and Cobolt 561 lasers and an additional 640 nm booster laser (iBeam Smart, Toptica) were collimated and focused onto a speckle reducer (catalog no. LSR-3005-17S-VIS, Optotune, Dietikon) before being coupled into a multi-mode fiber (catalog no. M105L02S-A, Thorlabs). The output of the fiber was magnified by an achromatic lens and focused into the sample to homogeneously illuminate an area of about 1,000 μm^2^. The laser was guided through a laser cleanup filter (390/482/563/640 HC Quad, AHF) to remove fluorescence generated by the fiber. Emitted fluorescence was collected through a high numerical aperture (NA) oil immersion objective (HCX PL APO 160×/1.43 NA, Leica), filtered by a 700/100 bandpass filter (catalog no. ET700/100m, Chroma) and imaged onto an Evolve512D EMCCD camera (Photometrics). For the filter settings for dual-color imaging see below (Ratiometric dual-color SMLM). The z focus was stabilized by an infrared laser that was totally internally reflected off the coverslip onto a quadrant photodiode, which was coupled into closed-loop feedback with the piezo objective positioner (Physik Instrumente). Laser control, focus stabilization and movement of filters was performed using a field-programmable gate array (Mojo, Embedded Micro). The pulse length of the 405 nm (laser intensity ≈ 28 W cm^-2^) laser was controlled by a feedback algorithm to sustain a predefined number of localizations per frame. Typical acquisition parameters are ~100,000 frames, a frame rate of 100 ms, and the laser intensity of 6 kW/cm^2^ as a good compromise between localization precision and imaging time^26^. Samples were mounted and imaged until almost all fluorophores were bleached and no further localizations were detected under continuous ultraviolet irradiation.

The yeast dual-color data were acquired on a microscope with a commercial laser box (iChrome MLE, Toptica, Gräfelfing) with 405 nm, 561 nm, and 640 nm lasers and a 640 nm booster laser (Toptica) which were coupled via single-mode. The output of the fiber was collimated, focused on the BFP of the TIRF objective (60× NA 1.49, Nikon), and adjusted for epi illumination. The emitted fluorescence was laterally constricted by a slit, split by a dichroic mirror (640LP, ZT640rdc, Chroma), filtered by the respective bandpass filters (transmitted/AF647: 676/37, FF01-676/37-25, Semrock; reflected/mMaple: 600/60, NC458462, Chroma), and imaged on two parts of the EMCCD camera (iXON Ultra, Andor). The focus was stabilized as described for the system above.

The microscopes were controlled by μManager^46^ through the Easier Micro-Manager User interface (EMU^47^).

#### Pixel size calibration

The effective pixel size of the microscope was calibrated by translating fluorescent beads, immobilized on a coverslip, with a calibrated sample stage (SmarAct) that operated in close loop. From the measured translation of many beads the pixel size could be calibrated with a high accuracy.

#### Ratiometric dual-color SMLM

For ratiometric dual-color imaging of AF647 and CF680, the emitted fluorescence was split by a 665LP beamsplitter (catalog no. ET665lp, Chroma), filtered by a 685/70 (catalog no. ET685/70m, Chroma) bandpass filter (transmitted light) or a 676/37 (catalog no. FF01-676/37-25, Semrock) bandpass filter (reflected light) and imaged side by side on the EMCCD camera. The color of the individual blinks was assigned by calculating the ratio of the intensities in the two channels.

#### Astigmatic 3D SMLM

The 3D SMLM data was acquired using a cylindrical lens (f= 1,000 mm; catalog no. LJ1516L1-A, Thorlabs) to introduce astigmatism. The data were fitted and analyzed as described previously^39^. First, *z* stacks with known displacement of several (15–20) fields of view of TetraSpeck beads on a coverslip were acquired to generate a model of the experimental point spread function. This model was then used to determine the z position of the individual localizations.

#### Dual-color SMLM in yeast

Raw data were acquired with 30 ms exposure time. The images acquired in the two channels (described above) were merged using a transformation which was determined using images of beads that are fluorescent in both channels (TetraSpeck).

### Data processing

SMLM data analysis was conducted using previously published algorithms with custom software written in MATLAB (super-resolution microscopy analysis platform, SMAP^21^), available as open source at github.com/jries/SMAP.

#### 3D bead calibration

Tetra-Speck beads (0.75 μL from stock, catalog no. T7279, Thermo Fisher) were diluted in 360 μL dH2O, mixed with 40 μL 1 M MgCl2 and put on a coverslip in a custom-manufactured sample holder. After 10 min, the mix was replaced with 400 μL dH2O. Using Micro-Manager, about 20 positions on the coverslip were defined and the beads were imaged acquiring *z* stacks (−1 to 1 μm, 10 nm step size) using the same filters as used in the intended experiment.

#### Fitting and postprocessing

Two-dimensional data were fitted with a symmetric Gaussian PSF model with the PSF size, *x, y*, photons per localization and the background as free-fitting parameters using maximum likelihood estimation^39^. 3D data were fitted using an experimentally derived PSF model from the 3D bead calibration with *x, y, z*, photons per localization and the background as free-fitting parameters using maximum likelihood estimation^39^.

Fitted data were first grouped by merging localizations persistent over consecutive frames within 35 nm from each other (with an allowed gap of one dark frame) into one localization with its position calculated by the weighted average of individual *x, y* and *z* positions. Photons per localization as well as the background were summed over all frames in which the grouped localization was detected. Data was then drift corrected in *x, y* and *z* by a custom algorithm based on redundant cross-correlation. From the spread of the redundant displacements we estimated the accuracy of the drift correction to be better than 1.5 nm in *x* and *y* and 2 nm in *z*.

To exclude bad fits and *z* position to reject molecules far away from the focal plane, the filtering was applied as follows.

3D data of Nup96 were filtered based on lateral localization precision ([0, 5] nm), *z* position (boundaries defined to exclude localizations away from the nuclear envelope), log-likelihood (lower boundary defined to exclude the left tail of the distribution), and frames (boundaries defined to exclude ~1,000 very first and last frames).

3D data of Microtubules were filtered based on lateral localization precision: ([0, 5] nm) and frames ([30,000, 90,000], also for efficiency).

3D dual-color data of NPCs were filtered based on lateral localization precision ([0, 10] nm for Nup96 and [0, 5] nm for target proteins), log-likelihood (lower boundary defined to exclude the left tail of the distribution), and frames (boundaries defined to exclude very first and last thousands of frames).

2D dual-color data of endocytic sites in yeast were filtered based on localization precision ([0, 25] nm), PSF size ([0, 175] nm), frames (boundaries defined to exclude ~20,000 very first and last frames).

#### Segmentation of sites

All NPC images used in this work that are based on Nup96-derived data, were segmented automatically in SMAP. For this, reconstructed images were convolved with a kernel consisting of a ring with a radius corresponding to the radius of the NPC, convolved with a Gaussian. Local maxima over a user-defined threshold were treated as possible candidates. Candidates were cleaned up by three additional steps. (1) We fitted the localizations corresponding to each candidate with a circle and excluded structures with a ring radius smaller than 40 nm or larger than 70 nm. (2) Localizations were refitted with a circle of fixed radius to determine the center coordinates. Structures were rejected if more than 25% of localizations were closer than 40 nm to the center or if more than 40% of localizations were further away than 70 nm from the center, as these typically did not visually resemble NPCs or were two adjacent wrongly segmented NPCs. (3) Sites with the number of localization smaller than 30 were further removed to ensure the sufficient sampling of the underlying biological structure.

In images of microtubules, a circular boundary with a radius of 500 nm was used to crop microtubules into sites to get segments that were at least one micrometer long. In a site with more than one microtubule, a polygon mask was used to further retain only one segment of interest.

Endocytic sites in yeast were manually picked and rotated so that the direction of the invagination was pointing upwards.

#### Correction of depth-dependent distortion

We observed a depth-dependent distortion along the *z*-axis, as reported previously^27^. The distortion is reflected by the depth-dependent ring separations *s* of NPCs (Extended Data Figure 4a). As the expected ring separation for Nup96 is known, we used it as the standard to correct the distortion. By definition the ring separation is the distance between the two rings of one NPC so that 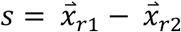, where 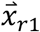 and 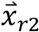 are the center positions of the two rings. As the orientation of an NPC is not necessary perpendicular to the *x-y* plane, we measured the tilt angle radian *ψ* of an NPC from the *z*-axis, and used the angle to derive the vertical component of the separation as *s_z_* = *s* cos *ψ* = *z*_*r*1_ – *z*_*r*2_. We also calculated the expected *s_z_* as E(*s_z_*) = E(*s*) cos*ψ*, with E(*s*) defined as 49.3 nm, the known average NPC ring separation^17^. With these values, we can calculate for each NPC a scaling factor *s_f_*(*z*) = E(*s_z_*)/*s_z_*. We found that the moving median of *s_z_* along z-axis appeared as a quadradic-like curve. We then fitted a quadradic function *s_z_* = *c*_2_*z*^2^ + *c*_2_*z* + *c*_3_ to the data. Given that the correction factor represents the change of the expected *z* position over the change of measured *z* position, *s_f_*(*z*) ≈ *∂E*(*z*)/*∂z*. We then defined *z*_0_, which makes *s_f_*(*z*) = 1, as the origin of distortion. The expected or undistorted *z* position can then be acquired as *E*(*z*) = ∫ *s_f_*(*z*)*dz* with *E*(*z*_0_) = *z*_0_. The corrected *z* position of each localization *k* was then defined as *z*_k_′ = *E*(*z_k_*) – *E*(0) for keeping the focal point zero. This correction was applied to all the NPC data sets before further quantifications.

## Acknowledgements

We thank I. Schoen and J. Hériché for inputs on the manuscript. This work was supported by the European Research Council (grant no. ERC CoG-724489 to J.R.), the National Institutes of Health Common Fund 4D Nucleome Program (grant no. U01 EB021223 to J.R.), the Human Frontier Science Program (grant no. RGY0065/2017 to J.R.), and the European Molecular Biology Laboratory.

## Author contributions

J.R. and Y.W. conceived the approach, developed the methods and wrote the software. U.M., P.H., A.T., and M.M. acquired the data. Y.W., A.T., P.H., M.M., and U.M. analyzed the data. Y.W. and J.R. wrote the manuscript with input from all authors.

## Extended Data Figures

**Extended Data Figure 1.**
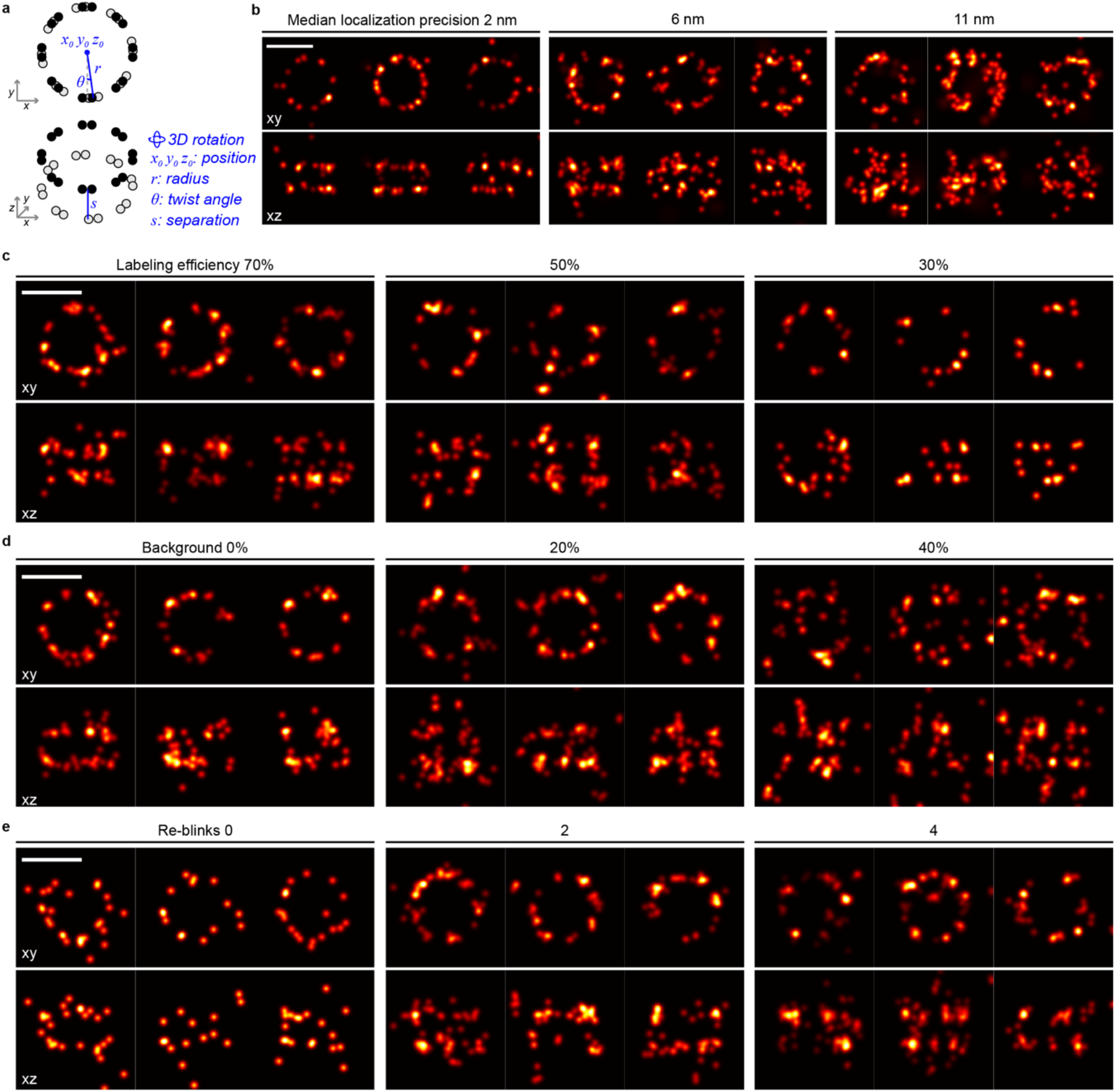
Example sites simulated across various conditions. **a,** the NPC model used for simulations. **b-e**, example simulated NPCs across different localization precision (**b**), labeling efficiency (**c**), background (**d**), and re-blinks (**e**). In **b**, corresponding photon counts *n_ph_* with zero fluorescence background *bg* were used to derive the specified median localization precisions. In **c-e**, all sites were simulated with *n_ph_* = 5,000 and *bg* = 100 photons/pixel/localization and yielded a median lateral localization precision of 3.6 nm. STORM mode and the condition as for **b** except for the parameters specified in the headers. Detailed simulation parameters are shown in Supplementary Table 1. Scale bars: 100 nm.

**Extended Data Figure 2.**
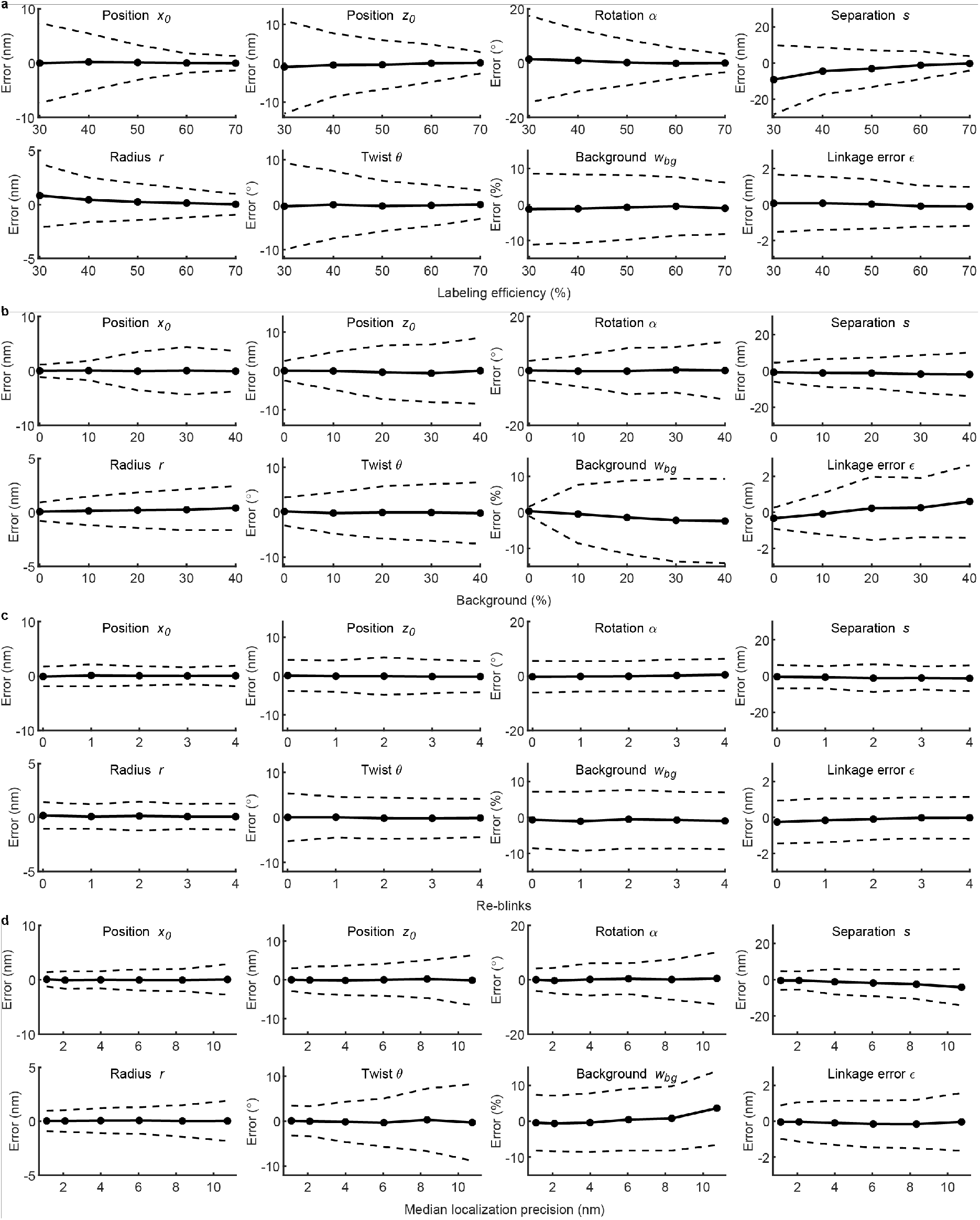
Errors of parameter estimations against factors contributing to data quality in the simulated datasets. Each panel shows the estimation errors of one parameter in the three imaging modes, across different levels of either labeling efficiency (a), background (b), re-blinks (c), or localization precision (d). Example sites of some conditions are shown in Extended Data Figure 1. The dots indicate mean errors and their standard deviations are indicated by dashed lines. Simulation parameters and plotted statistics are shown in Supplementary Table 1 and Supplementary Table 2, respectively. Sample size: ns = 1,000 for each dot.

**Extended Data Figure 3.**
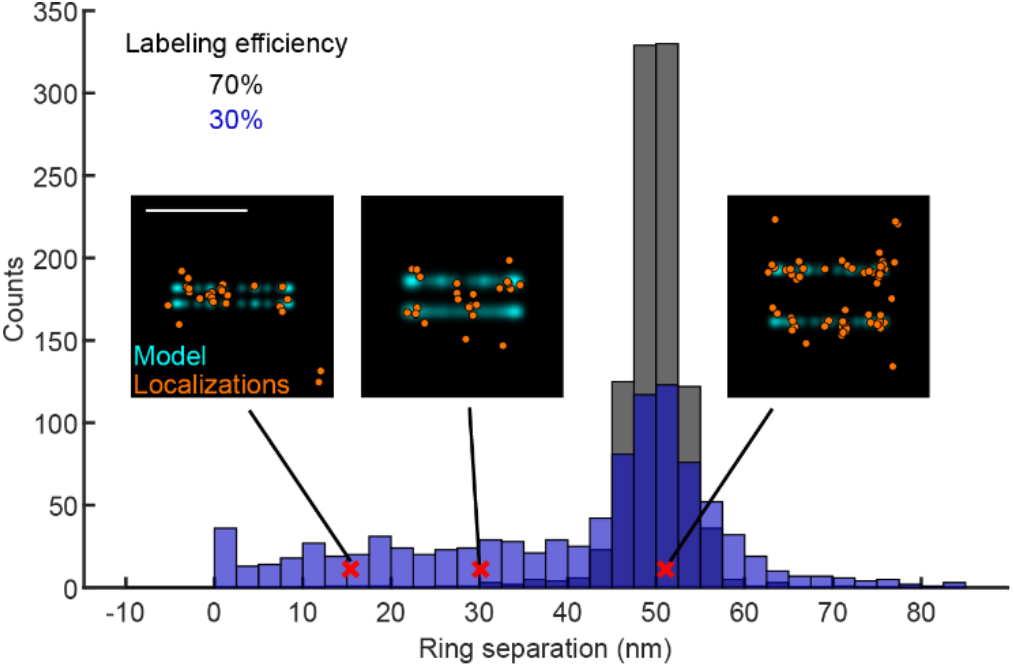
Distributions of the fitted ring separation at different labeling efficiencies in simulated data. Three side-view examples of pores at different fitted ring separations, indicated by red crosses. All examples are from the dataset simulated with 30% labeling efficiency. Scale bar: 100 nm.

**Extended Data Figure 4.**
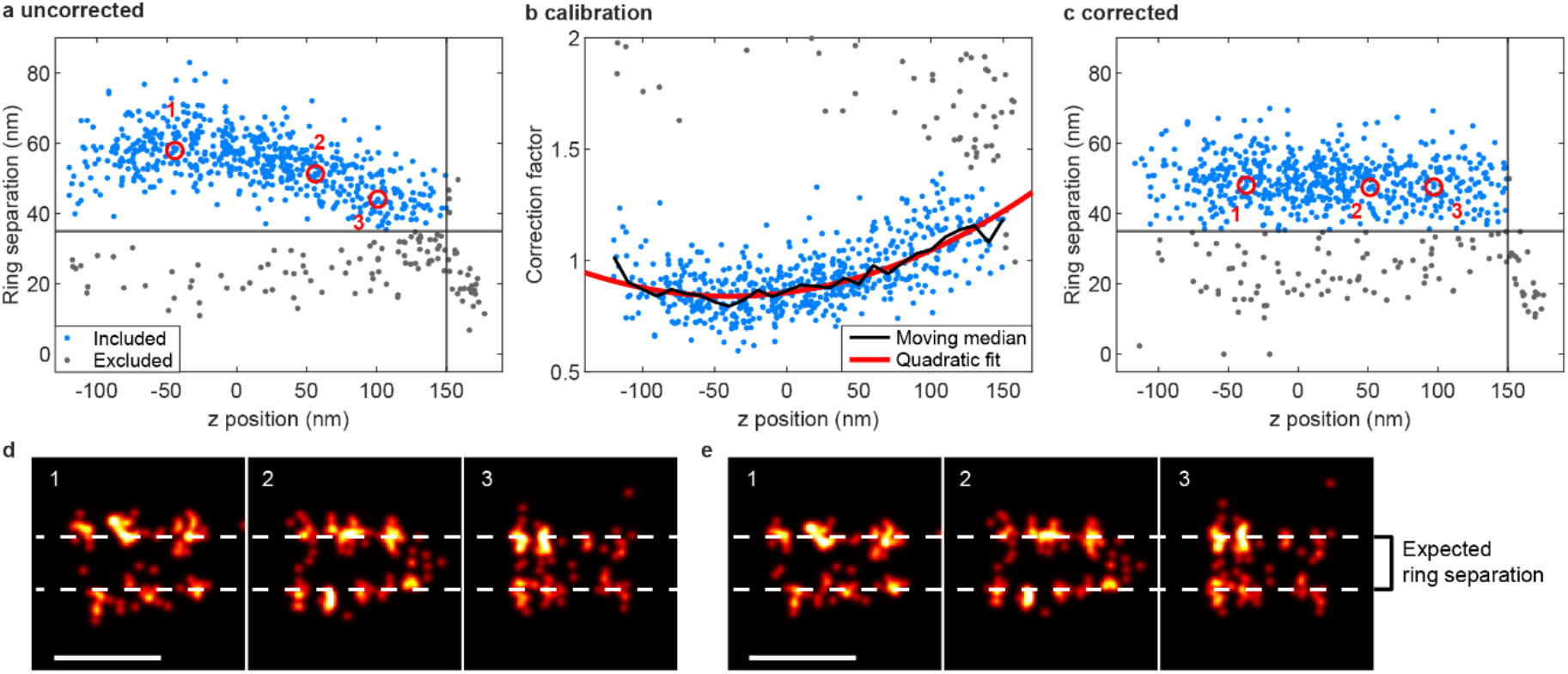
Correction of *z*-dependent distortion. **a**. A scatter plot showing the *z*-dependent spatial distortion along the *z* axis, as the NPC ring separation appears dependent on the *z* position. **b**. The median correction factor over *z* positions can be approximated by a quadratic fit. The correction factor is defined as the expected ring separation divided by a fitted ring separation before the correction. **c**. Fitted ring separations over the *z* positions after the correction based on the correction factor. NPCs with ring separations smaller than 35 nm (gray horizontal lines, likely stemming from NPCs with only a single ring labeled) and with *z* positions further than 150 nm away from the focus (gray vertical lines) were excluded from the curve fit and the following analysis. Each point represents one NPC. All the data points in a-c are from the same field of view. Side-view examples of sites at different *z* positions before (**d**) and after (**e**) the correction, corresponding to the data points indicated by the numbered red circles in **a** and **c**. Scale bars: 100 nm.

**Extended Data Figure 5.**
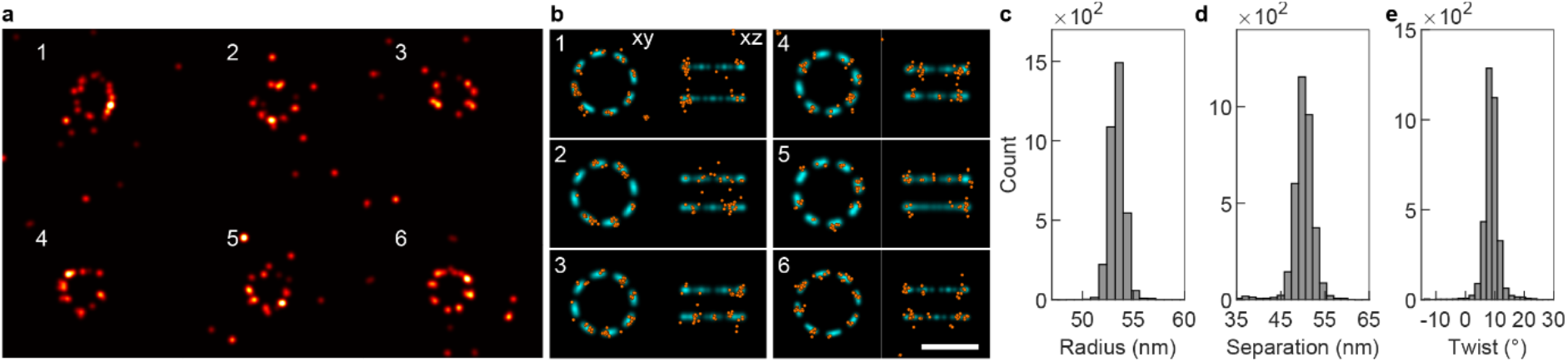
Simulations based on experimental parameters. **a**, Representative images of the simulated Nup96-labeled NPCs in 3D (top view). **b**, single NPCs (orange) as indicated in **a** are fitted with the eight-fold symmetric model. **c-e**, Histograms of three fitted parameters: radius r = 53.5 ± 1.5 nm (**c**), separation s = 50.3 ± 3.8 nm (**d**), and twist θ = 8.6 ± 5.9° (**e**). Simulation parameters are summarized in Supplementary Table 1. Sample size: n_s_ = 4,000. Scale bars: 100 nm.

**Extended Data Figure 6.**
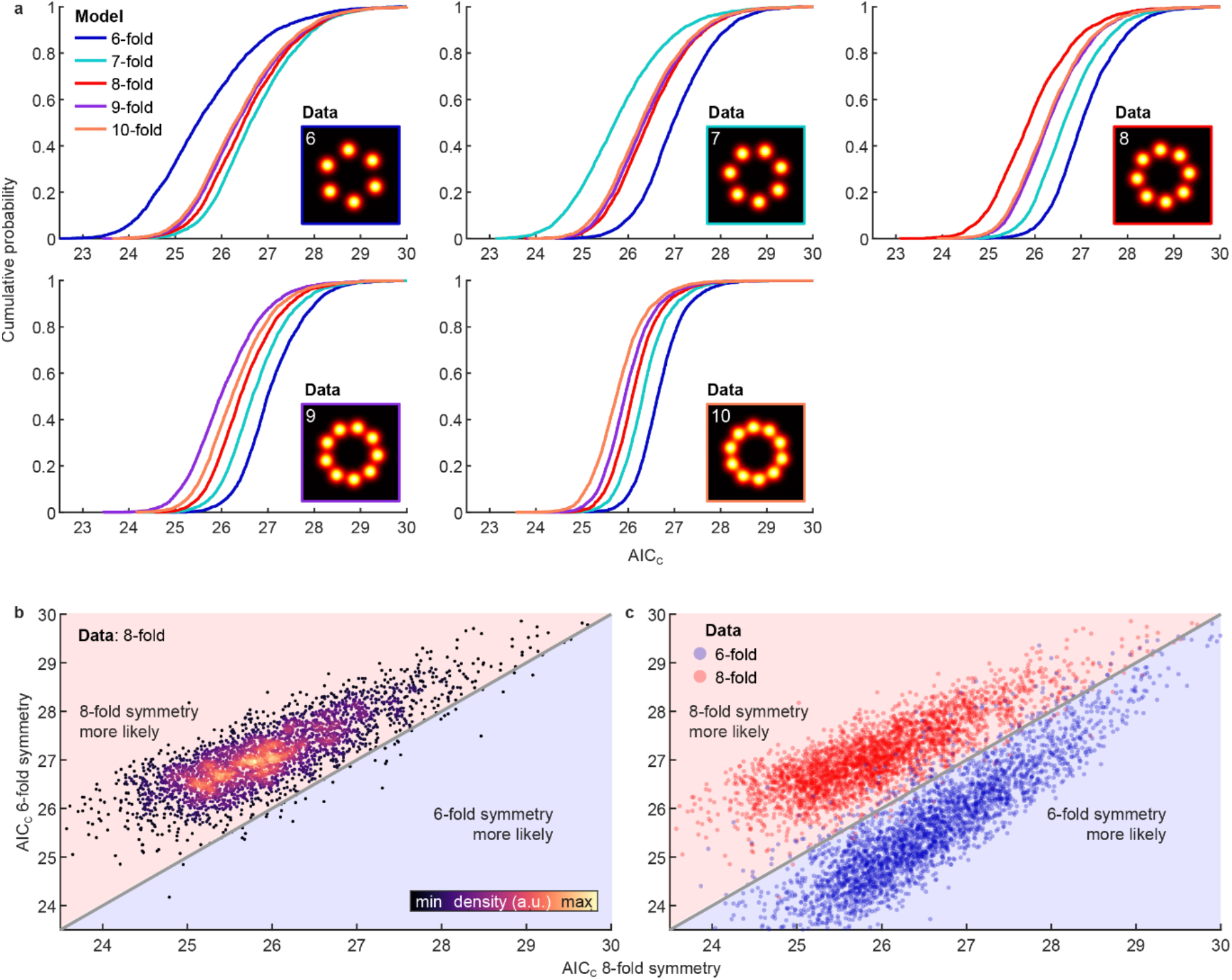
Validation of the model selection using simulations. **a**, Normalized AIC_C_ of fitting simulated NPCs with different rotationally symmetric models. Each panel corresponds to a simulated NPC dataset generated from the model indicated in the insets. Sample size: *n*_s_ = 3,000 for each panel. **b**, Scatter plot showing normalized AIC_C_ values of fitting the simulated 8-fold symmetric sites with models having 6-fold and 8-fold rotational symmetries. **c**, Scatter plot showing normalized AIC_C_ values of fitting both the simulated six-fold and eight-fold symmetric sites with models having six-fold and eight-fold rotational symmetries. The gray diagonal lines indicate equal normalized AIC_C_ values. Simulation parameters are summarized in Supplementary Table 1 and correspond to SMLM in the dSTORM mode.

## Supplementary information

**Supplementary Table 1.**
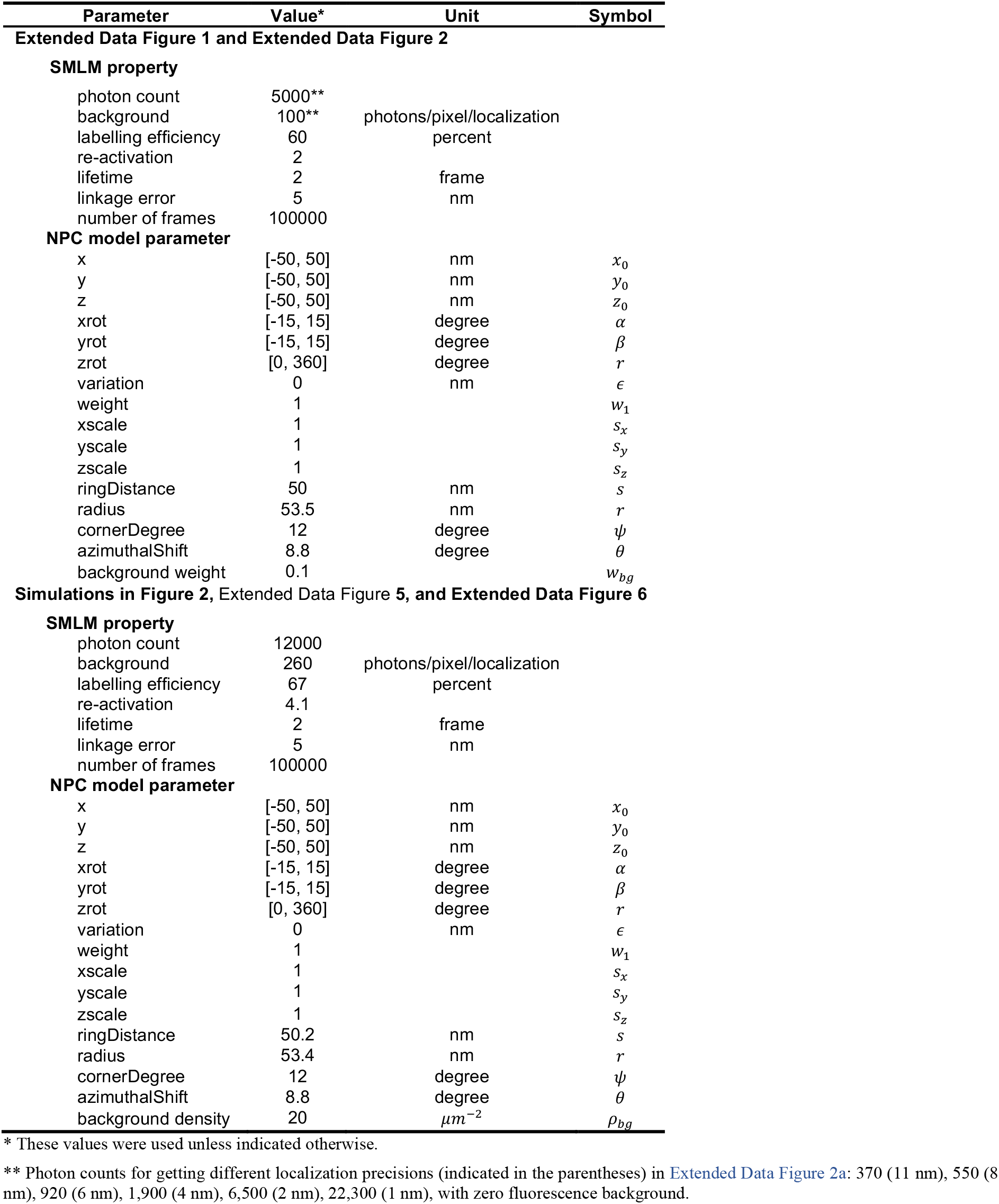
Simulation parameters.

**Supplementary Table 2.** Data points plotted in Extended Data Figure 2

| Parameter | Data point* | mean error ±<br>standard deviation | Unit |
| --- | --- | --- | --- |
| $x_0$ | locprec_1 | $0.1 \pm 1.3$ | nm |
| $x_0$ | locprec_2 | $-0.1 \pm 1.6$ | nm |
| $x_0$ | locprec_4 | $0.0 \pm 1.6$ | nm |
| $x_0$ | locprec_6 | $0.0 \pm 1.9$ | nm |
| $x_0$ | locprec_8 | $0.0 \pm 2.0$ | nm |
| $x_0$ | locprec_11 | $0.1 \pm 2.8$ | nm |
| $z_0$ | locprec_1 | $0.0 \pm 2.9$ | nm |
| $z_0$ | locprec_2 | $-0.1 \pm 3.4$ | nm |
| $z_0$ | locprec_4 | $-0.2 \pm 3.8$ | nm |
| $z_0$ | locprec_6 | $0.0 \pm 4.1$ | nm |
| $z_0$ | locprec_8 | $0.2 \pm 4.9$ | nm |
| $z_0$ | locprec_11 | $-0.1 \pm 6.4$ | nm |
| $\alpha$ | locprec_1 | $0.0 \pm 4.1$ | ° |
| $\alpha$ | locprec_2 | $-0.3 \pm 4.7$ | ° |
| $\alpha$ | locprec_4 | $0.1 \pm 5.9$ | ° |
| $\alpha$ | locprec_6 | $0.4 \pm 5.6$ | ° |
| $\alpha$ | locprec_8 | $0.1 \pm 7.4$ | ° |
| $\alpha$ | locprec_11 | $0.5 \pm 9.5$ | ° |
| $s$ | locprec_1 | $-0.4 \pm 5.2$ | nm |
| $s$ | locprec_2 | $-0.4 \pm 5.0$ | nm |
| $s$ | locprec_4 | $-1.1 \pm 7.1$ | nm |
| $s$ | locprec_6 | $-1.8 \pm 7.3$ | nm |
| $s$ | locprec_8 | $-2.5 \pm 8.1$ | nm |
| $s$ | locprec_11 | $-4.1 \pm 10.1$ | nm |
| $r$ | locprec_1 | $0.0 \pm 1.0$ | nm |
| $r$ | locprec_2 | $0.0 \pm 1.0$ | nm |
| $r$ | locprec_4 | $0.1 \pm 1.2$ | nm |
| $r$ | locprec_6 | $0.1 \pm 1.2$ | nm |
| $r$ | locprec_8 | $0.0 \pm 1.5$ | nm |
| $r$ | locprec_11 | $0.0 \pm 1.9$ | nm |
| $\theta$ | locprec_1 | $0.1 \pm 3.3$ | ° |
| $\theta$ | locprec_2 | $0.0 \pm 3.3$ | ° |
| $\theta$ | locprec_4 | $-0.1 \pm 4.5$ | ° |
| $\theta$ | locprec_6 | $-0.3 \pm 5.4$ | ° |
| $\theta$ | locprec_8 | $0.3 \pm 7.0$ | ° |
| $\theta$ | locprec_11 | $-0.3 \pm 8.5$ | ° |
| $w_{bg}$ | locprec_1 | $-0.4 \pm 7.8$ | % |
| $w_{bg}$ | locprec_2 | $-0.6 \pm 7.7$ | % |
| $w_{bg}$ | locprec_4 | $-0.4 \pm 8.1$ | % |
| $w_{bg}$ | locprec_6 | $0.5 \pm 8.6$ | % |
| $w_{bg}$ | locprec_8 | $0.8 \pm 8.9$ | % |
| $w_{bg}$ | locprec_11 | $3.6 \pm 10.3$ | % |
| $\epsilon$ | locprec_1 | $0.0 \pm 0.9$ | nm |
| $\epsilon$ | locprec_2 | $0.0 \pm 1.1$ | nm |
| $\epsilon$ | locprec_4 | $-0.1 \pm 1.2$ | nm |
| $\epsilon$ | locprec_6 | $-0.2 \pm 1.3$ | nm |
| $\epsilon$ | locprec_8 | $-0.2 \pm 1.3$ | nm |
| $\epsilon$ | locprec_11 | $0.0 \pm 1.6$ | nm |
| $x_0$ | bg_0 | $0.0 \pm 1.1$ | nm |
| $x_0$ | bg_10 | $0.0 \pm 1.8$ | nm |
| $x_0$ | bg_20 | $0.0 \pm 3.5$ | nm |
| $x_0$ | bg_30 | $0.0 \pm 4.4$ | nm |
| $x_0$ | bg_40 | $-0.1 \pm 3.7$ | nm |
| $z_0$ | bg_0 | $0.0 \pm 2.5$ | nm |
| $z_0$ | bg_10 | $0.0 \pm 4.8$ | nm |
| $z_0$ | bg_20 | $-0.4 \pm 6.9$ | nm |
| $z_0$ | bg_30 | $-0.6 \pm 7.4$ | nm |
| $z_0$ | bg_40 | $0.0 \pm 8.5$ | nm |
| $\alpha$ | bg_0 | $0.1 \pm 3.6$ | ° |
| $\alpha$ | bg_10 | $-0.1 \pm 5.6$ | ° |
| $\alpha$ | bg_20 | $-0.1 \pm 8.4$ | ° |
| $\alpha$ | bg_30 | $0.4 \pm 8.4$ | ° |
| $\alpha$ | bg_40 | $0.1 \pm 10.6$ | ° |
| $s$ | bg_0 | $-0.8 \pm 5.2$ | nm |
| $s$ | bg_10 | $-1.1 \pm 7.7$ | nm |
| $s$ | bg_20 | $-1.3 \pm 8.5$ | nm |
| $s$ | bg_30 | $-1.8 \pm 10.4$ | nm |
| $s$ | bg_40 | $-1.9 \pm 11.9$ | nm |
| $r$ | bg_0 | $0.1 \pm 0.8$ | nm |
| $r$ | bg_10 | $0.1 \pm 1.3$ | nm |
| $r$ | bg_20 | $0.2 \pm 1.6$ | nm |
| $r$ | bg_30 | $0.2 \pm 1.9$ | nm |
| $r$ | bg_40 | $0.4 \pm 2.0$ | nm |
| $\theta$ | bg_0 | $0.2 \pm 3.1$ | ° |
| $\theta$ | bg_10 | $-0.2 \pm 4.6$ | ° |
| $\theta$ | bg_20 | $0.0 \pm 5.8$ | ° |
| $\theta$ | bg_30 | $0.0 \pm 6.3$ | ° |
| $\theta$ | bg_40 | $-0.2 \pm 6.8$ | ° |
| $w_{bg}$ | bg_0 | $0.3 \pm 1.3$ | % |
| $w_{bg}$ | bg_10 | $-0.5 \pm 8.1$ | % |
| $w_{bg}$ | bg_20 | $-1.4 \pm 10.2$ | % |
| $w_{bg}$ | bg_30 | $-2.2 \pm 11.5$ | % |
| $w_{bg}$ | bg_40 | $-2.4 \pm 11.7$ | % |
| $\epsilon$ | bg_0 | $-0.3 \pm 0.6$ | nm |
| $\epsilon$ | bg_10 | $-0.1 \pm 1.1$ | nm |
| $\epsilon$ | bg_20 | $0.2 \pm 1.8$ | nm |
| $\epsilon$ | bg_30 | $0.3 \pm 1.6$ | nm |
| $\epsilon$ | bg_40 | $0.6 \pm 2.0$ | nm |
| $x_0$ | LE_70 | $0.0 \pm 1.3$ | nm |
| $x_0$ | LE_60 | $0.0 \pm 1.8$ | nm |
| $x_0$ | LE_50 | $0.1 \pm 3.2$ | nm |
| $x_0$ | LE_40 | $0.2 \pm 5.2$ | nm |
| $x_0$ | LE_30 | $0.0 \pm 7.4$ | nm |
| $z_0$ | LE_70 | $0.1 \pm 2.8$ | nm |
| $z_0$ | LE_60 | $0.0 \pm 4.8$ | nm |
| $z_0$ | LE_50 | $-0.4 \pm 6.3$ | nm |
| $z_0$ | LE_40 | $-0.5 \pm 8.2$ | nm |
| $z_0$ | LE_30 | $-1.0 \pm 12.0$ | nm |
| $\alpha$ | LE_70 | $0.1 \pm 3.4$ | ° |
| $\alpha$ | LE_60 | $-0.1 \pm 5.6$ | ° |
| $\alpha$ | LE_50 | $0.2 \pm 8.4$ | ° |
| $\alpha$ | LE_40 | $1.0 \pm 11.4$ | ° |
| $\alpha$ | LE_30 | $1.5 \pm 16.0$ | ° |
| $s$ | LE_70 | $-0.2 \pm 4.0$ | nm |
| $s$ | LE_60 | $-1.1 \pm 7.7$ | nm |
| $s$ | LE_50 | $-3.0 \pm 10.2$ | nm |
| $s$ | LE_40 | $-4.4 \pm 13.0$ | nm |
| $s$ | LE_30 | $-9.1 \pm 19.1$ | nm |
| $r$ | LE_70 | $0.0 \pm 1.0$ | nm |
| $r$ | LE_60 | $0.1 \pm 1.3$ | nm |
| $r$ | LE_50 | $0.2 \pm 1.7$ | nm |
| $r$ | LE_40 | $0.4 \pm 2.1$ | nm |
| $r$ | LE_30 | $0.8 \pm 3.0$ | nm |
| $\theta$ | LE_70 | $0.0 \pm 3.2$ | ° |
| $\theta$ | LE_60 | $-0.2 \pm 4.6$ | $^{\circ}$ |
| $\theta$ | LE_50 | $-0.3 \pm 5.6$ | $^{\circ}$ |
| $\theta$ | LE_40 | $0.0 \pm 7.5$ | $^{\circ}$ |
| $\theta$ | LE_30 | $-0.4 \pm 9.6$ | $^{\circ}$ |
| $w_{bg}$ | LE_70 | $-1.0 \pm 7.2$ | % |
| $w_{bg}$ | LE_60 | $-0.5 \pm 8.1$ | % |
| $w_{bg}$ | LE_50 | $-0.8 \pm 9.0$ | % |
| $w_{bg}$ | LE_40 | $-1.1 \pm 9.5$ | % |
| $w_{bg}$ | LE_30 | $-1.3 \pm 9.9$ | % |
| $\epsilon$ | LE_70 | $-0.1 \pm 1.1$ | nm |
| $\epsilon$ | LE_60 | $-0.1 \pm 1.1$ | nm |
| $\epsilon$ | LE_50 | $0.0 \pm 1.4$ | nm |
| $\epsilon$ | LE_40 | $0.1 \pm 1.5$ | nm |
| $\epsilon$ | LE_30 | $0.1 \pm 1.6$ | nm |
| $x_0$ | RB_0 | $-0.1 \pm 1.8$ | nm |
| $x_0$ | RB_1 | $0.1 \pm 2.0$ | nm |
| $x_0$ | RB_2 | $0.0 \pm 1.8$ | nm |
| $x_0$ | RB_3 | $0.0 \pm 1.6$ | nm |
| $x_0$ | RB_4 | $0.0 \pm 1.9$ | nm |
| $z_0$ | RB_0 | $0.1 \pm 4.0$ | nm |
| $z_0$ | RB_1 | $0.0 \pm 4.0$ | nm |
| $z_0$ | RB_2 | $0.0 \pm 4.8$ | nm |
| $z_0$ | RB_3 | $-0.1 \pm 4.3$ | nm |
| $z_0$ | RB_4 | $-0.1 \pm 4.0$ | nm |
| $\alpha$ | RB_0 | $-0.3 \pm 5.9$ | $^{\circ}$ |
| $\alpha$ | RB_1 | $-0.1 \pm 5.6$ | $^{\circ}$ |
| $\alpha$ | RB_2 | $-0.1 \pm 5.6$ | $^{\circ}$ |
| $\alpha$ | RB_3 | $0.2 \pm 5.9$ | $^{\circ}$ |
| $\alpha$ | RB_4 | $0.5 \pm 5.9$ | $^{\circ}$ |
| $s$ | RB_0 | $-0.4 \pm 6.4$ | nm |
| $s$ | RB_1 | $-0.7 \pm 6.2$ | nm |
| $s$ | RB_2 | $-1.1 \pm 7.7$ | nm |
| $s$ | RB_3 | $-1.1 \pm 6.4$ | nm |
| $s$ | RB_4 | $-1.3 \pm 7.1$ | nm |
| $r$ | RB_0 | $0.2 \pm 1.2$ | nm |
| $r$ | RB_1 | $0.1 \pm 1.1$ | nm |
| $r$ | RB_2 | $0.1 \pm 1.3$ | nm |
| $r$ | RB_3 | $0.1 \pm 1.2$ | nm |
| $r$ | RB_4 | $0.1 \pm 1.2$ | nm |
| $\theta$ | RB_0 | $0.0 \pm 5.3$ | $^{\circ}$ |
| $\theta$ | RB_1 | $0.1 \pm 4.5$ | $^{\circ}$ |
| $\theta$ | RB_2 | $-0.2 \pm 4.6$ | $^{\circ}$ |
| $\theta$ | RB_3 | $-0.2 \pm 4.4$ | $^{\circ}$ |
| $\theta$ | RB_4 | $-0.1 \pm 4.3$ | $^{\circ}$ |
| $w_{bg}$ | RB_0 | $-0.7 \pm 7.8$ | % |
| $w_{bg}$ | RB_1 | $-1.1 \pm 8.3$ | % |
| $w_{bg}$ | RB_2 | $-0.5 \pm 8.1$ | % |
| $w_{bg}$ | RB_3 | $-0.7 \pm 7.9$ | % |
| $w_{bg}$ | RB_4 | $-1.0 \pm 7.9$ | % |
| $\epsilon$ | RB_0 | $-0.3 \pm 1.2$ | nm |
| $\epsilon$ | RB_1 | $-0.2 \pm 1.2$ | nm |
| $\epsilon$ | RB_2 | $-0.1 \pm 1.1$ | nm |
| $\epsilon$ | RB_3 | $0.0 \pm 1.1$ | nm |
| $\epsilon$ | RB_4 | $0.0 \pm 1.2$ | nm |

\* Abbreviations: ‘locprec’ for localization precision, ‘LE’ for labeling efficiency, ‘BG’ for background, and ‘RB’ for re-blink.

**Supplementary Table 3.** Fitting settings used in this study for Nup96 in single color data.

| Step | Model | Parameter type | Internal parameter name | Free parameter | Relative |  | Absolute |  | Initial value | Unit | Symbol |
| --- | --- | --- | --- | --- | --- | --- | --- | --- | --- | --- | --- |
|  |  |  |  |  | LB | UB | LB | UB |  |  |  |
| 1 | 1 | e | x | yes | -50 | 50 | -150 | 150 | $\tilde{x}_k$ | nm | $x_0$ |
| 1 | 1 | e | y | yes | -50 | 50 | -150 | 150 | $\tilde{y}_k$ | nm | $y_0$ |
| 1 | 1 | e | z | yes | -100 | 20 | -300 | 300 | $\tilde{z}_k - 40$ | nm | $z_0$ |
| 1 | 1 | e | weight | no | | | | | 1 | | $w_1$ |
| 1 | 1 | e | xrot | yes | -30 | 30 | -inf | inf | 0 | ° | $\alpha$ |
| 1 | 1 | e | yrot | yes | -30 | 30 | -inf | inf | 0 | ° | $\beta$ |
| 1 | 1 | e | zrot | no | | | | | 0 | ° | $\gamma$ |
| 1 | 1 | e | xscale | no | | | | | 1 | | $s_x$ |
| 1 | 1 | e | yscale | no | | | | | 1 | | $s_y$ |
| 1 | 1 | e | zscale | no | | | | | 1 | | $s_z$ |
| 1 | 1 | e | variation | no | | | | | 0 | nm | $\epsilon$ |
| 1 | 2 | e | x | no | | | | | 0 | nm | $x_0$ |
| 1 | 2 | e | y | no | | | | | 0 | nm | $y_0$ |
| 1 | 2 | e | z | yes | -40 | 100 | 0 | 300 | 40 | nm | $\hat{z}_0$ |
| 1 | 2 | e | xrot | no | | | | | 0 | ° | $\alpha$ |
| 1 | 2 | e | yrot | no | | | | | 0 | ° | $\beta$ |
| 1 | 2 | e | zrot | no | | | | | 0 | ° | $\gamma$ |
| 1 | 2 | e | variation | no | | | | | 0 | nm | $\epsilon$ |
| 1 | 2 | e | weight | no | | | | | 1 | | $w_1$ |
| 1 | 2 | e | xscale | no | | | | | 0 | | $s_x$ |
| 1 | 2 | e | yscale | no | | | | | 0 | | $s_y$ |
| 1 | 2 | e | zscale | no | | | | | 0 | | $s_z$ |
| 1 | L1 | e | weight | yes | -1 | 1 | 0.001 | 0.999 | 0 | | $w_{bg}$ |
| 2 | 1 | e | x | yes | -20 | 20 | -150 | 150 | $\hat{x}_0^{1,1} + \phi(\hat{z}_0^{1,2})_{[1]}$ | nm | $x_0$ |
| 2 | 1 | e | y | yes | -20 | 20 | -150 | 150 | $\hat{y}_0^{1,1} + \phi(\hat{z}_0^{1,2})_{[2]}$ | nm | $y_0$ |
| 2 | 1 | e | z | yes | -20 | 20 | -300 | 300 | $\hat{z}_0^{1,1} + \phi(\hat{z}_0^{1,2})_{[3]}$ | nm | $z_0$ |
| 2 | 1 | e | xrot | yes | -15 | 15 | -inf | inf | inherited | ° | $\alpha$ |
| 2 | 1 | e | yrot | yes | -15 | 15 | -inf | inf | inherited | ° | $\beta$ |
| 2 | 1 | e | zrot | no | | | | | 0 | ° | $\gamma$ |
| 2 | 1 | e | variation | yes | 0 | 20 | 1 | 20 | 0 | nm | $\epsilon$ |
| 2 | 1 | e | weight | no | | | | | 1 | | $w_1$ |
| 2 | 1 | e | xscale | no | | | | | 0 | | $s_x$ |
| 2 | 1 | e | yscale | no | | | | | 0 | | $s_y$ |
| 2 | 1 | e | zscale | no | | | | | 0 | | $s_z$ |
| 2 | 1 | i | azimuthalShift | no | | | | | 0 | ° | $\theta$ |
| 2 | 1 | i | cornerDegree | no | | | | | 12 | ° | $\psi$ |
| 2 | 1 | i | ringDistance | yes | -10 | 10 | 0 | 100 | inherited | nm | $s$ |
| 2 | 1 | i | radius | yes | -10 | 10 | 30 | 70 | inherited | nm | $r$ |
| 2 | L1 | e | weight | no | | | | | inherited | | $w_{bg}$ |
| 3 | 1 | e | x | no | | | | | inherited | nm | $x_0$ |
| 3 | 1 | e | y | no | | | | | inherited | nm | $y_0$ |
| 3 | 1 | e | z | no | | | | | inherited | nm | $z_0$ |
| 3 | 1 | e | xrot | no | | | | | inherited | ° | $\alpha$ |
| 3 | 1 | e | yrot | no | | | | | inherited | ° | $\beta$ |
| 3 | 1 | e | zrot | yes | -180 | 180 | -inf | inf | 0 | ° | $r$ |
| 3 | 1 | e | variation | yes | -10 | 10 | 1 | 20 | 5 | nm | $\epsilon$ |
| 3 | 1 | e | weight | no | | | | | inherited | | $w_1$ |
| 3 | 1 | e | xscale | no | | | | | inherited | | $s_x$ |
| 3 | 1 | e | yscale | no | | | | | inherited | | $s_y$ |
| 3 | 1 | e | zscale | no | | | | | inherited | | $s_z$ |
| 3 | 1 | i | ringDistance | no | | | | | inherited | nm | $s$ |
| 3 | 1 | i | radius | no | | | | | inherited | nm | $r$ |
| 3 | 1 | i | cornerDegree | no | | | | | 12 | ° | $\psi$ |
| 3 | 1 | i | azimuthalShift | yes | -180 | 180 | -inf | inf | 0 | ° | $\theta$ |
| 3 | L1 | e | weight | yes | -1 | 1 | 0.001 | 0.999 | inherited | | $w_{bg}$ |

Note: In column *Parameter type*, *i* and *e* represent intrinsic and extrinsic parameters, respectively. In column *Model*, L1 represents layer 1. In column *initial value*, a tilde operator represents a median value, a hat operator indicates a parameter estimate, and superscripts represent steps and models (e.g., $\hat{z}_0^{1,2}$ means the estimate of $z_0$ from model 2 in step 1). The function $\vec{z}'_{[k]} = \phi(z)_{[k]}$ outputs the $k^{\text{th}}$ element of the vector $\vec{z}'$ derived from $\vec{z} = [0 \ 0 \ z]$ rotated by the rotation matrix constructed using $\{\hat{\alpha}^{1,1}, \hat{\beta}^{1,1}, \hat{\gamma}^{1,1}\}$ as the angles $\{\alpha, \beta, \gamma\}$ in the equation
(11). ‘inherited’ indicates that a value is inherited from its counterpart estimate in the last step. Other symbols have the same meanings as in the main text. Abbreviations: LB/UB: lower/upper boundaries. Blanks indicate values not applicable.

**Supplementary Table 4.** Fitting settings used in this study for microtubules.

| Step | Model | Parameter type | Internal parameter name | Free parameter | Relative |  | Absolute |  | Initial value | Unit | Symbol |
| --- | --- | --- | --- | --- | --- | --- | --- | --- | --- | --- | --- |
|  |  |  |  |  | LB | UB | LB | UB |  |  |  |
| 1 | 1 | *e* | x | no | | | | | $\tilde{x}_k$ | nm | $x_0$ |
| 1 | 1 | *e* | y | no | | | | | $\tilde{y}_k$ | nm | $y_0$ |
| 1 | 1 | *e* | z | no | | | | | $\tilde{z}_k$ | nm | $z_0$ |
| 1 | 1 | *e* | xrot | no | | | | | 0 | ° | $\alpha$ |
| 1 | 1 | *e* | yrot | no | | | | | 0 | ° | $\beta$ |
| 1 | 1 | *e* | zrot | no | | | | | $pcaRot(\tilde{x}_k, \tilde{y}_k)$ | ° | $\gamma$ |
| 1 | 1 | *e* | variation | yes | -Inf | Inf | 30 | 45 | 40 | nm | $\epsilon$ |
| 1 | 1 | *e* | weight | no | | | | | 1 | | $w_1$ |
| 1 | 1 | *e* | xscale | no | | | | | 1 | | $s_x$ |
| 1 | 1 | *e* | yscale | no | | | | | 1 | | $s_y$ |
| 1 | 1 | *e* | zscale | no | | | | | 1 | | $s_z$ |
| 1 | 1 | *i* | xMid | yes | -Inf | Inf | -500 | 500 | 0 | nm | $x_q$ |
| 1 | 1 | *i* | yMid | yes | -Inf | Inf | -500 | 500 | 0 | nm | $y_q$ |
| 1 | 1 | *i* | zMid | yes | -Inf | Inf | -500 | 500 | 0 | nm | $z_q$ |
| 1 | 1 | *i* | dist | no | | | | | 250 | nm | $q$ |
| 1 | 1 | *i* | rotAziL1 | yes | -25 | 25 | -Inf | Inf | 0 | ° | $\theta_{L1}$ |
| 1 | 1 | *i* | rotEleL1 | yes | -25 | 25 | -Inf | Inf | 0 | ° | $\phi_{L1}$ |
| 1 | 1 | *i* | rotAziR1 | yes | -25 | 25 | -Inf | Inf | 0 | ° | $\theta_{R1}$ |
| 1 | 1 | *i* | rotEleR1 | yes | -25 | 25 | -Inf | Inf | 0 | ° | $\phi_{R1}$ |
| 1 | 1 | *i* | rotAziL2 | yes | -25 | 25 | -Inf | Inf | 0 | ° | $\theta_{L2}$ |
| 1 | 1 | *i* | rotEleL2 | yes | -25 | 25 | -Inf | Inf | 0 | ° | $\phi_{L2}$ |
| 1 | 1 | *i* | rotAziR2 | yes | -25 | 25 | -Inf | Inf | 0 | ° | $\theta_{R1}$ |
| 1 | 1 | *i* | rotEleR2 | yes | -25 | 25 | -Inf | Inf | 0 | ° | $\phi_{R1}$ |
| 1 | L1 | *e* | weight | yes | -1 | 1 | 0.001 | 0.999 | 0.5 | | $w_{bg}$ |
| 2 | 1 | *e* | x | no | | | | | inherited | nm | $x_0$ |
| 2 | 1 | *e* | y | no | | | | | inherited | nm | $y_0$ |
| 2 | 1 | *e* | z | no | | | | | inherited | nm | $z_0$ |
| 2 | 1 | *e* | xrot | no | | | | | inherited | ° | $\alpha$ |
| 2 | 1 | *e* | yrot | no | | | | | inherited | ° | $\beta$ |
| 2 | 1 | *e* | zrot | no | | | | | inherited | ° | $\gamma$ |
| 2 | 1 | *e* | variation | no | | | | | 7 | nm | $\epsilon$ |
| 2 | 1 | *e* | weight | no | | | | | inherited | | $w_1$ |
| 2 | 1 | *e* | xscale | no | | | | | inherited | | $s_x$ |
| 2 | 1 | *e* | yscale | no | | | | | inherited | | $s_y$ |
| 2 | 1 | *e* | zscale | no | | | | | inherited | | $s_z$ |
| 2 | 1 | *i* | xMid | yes | -30 | 30 | -500 | 500 | inherited | nm | $x_q$ |
| 2 | 1 | *i* | yMid | yes | -30 | 30 | -500 | 500 | inherited | nm | $y_q$ |
| 2 | 1 | *i* | zMid | yes | -30 | 30 | -500 | 500 | inherited | nm | $z_q$ |
| 2 | 1 | *i* | r | yes | -Inf | Inf | 10 | 60 | 30 | nm | $r$ |
| 2 | 1 | *i* | dist | no | | | | | inherited | nm | $q$ |
| 2 | 1 | *i* | rotAziL1 | yes | -Inf | Inf | -25 | 25 | inherited | ° | $\theta_{L1}$ |
| 2 | 1 | *i* | rotEleL1 | yes | -Inf | Inf | -25 | 25 | inherited | ° | $\phi_{L1}$ |
| 2 | 1 | *i* | rotAziR1 | yes | -Inf | Inf | -25 | 25 | inherited | ° | $\theta_{R1}$ |
| 2 | 1 | *i* | rotEleR1 | yes | -Inf | Inf | -25 | 25 | inherited | ° | $\phi_{R1}$ |
| 2 | 1 | *i* | rotAziL2 | yes | -Inf | Inf | -25 | 25 | inherited | ° | $\theta_{L2}$ |
| 2 | 1 | *i* | rotEleL2 | yes | -Inf | Inf | -25 | 25 | inherited | ° | $\phi_{L2}$ |
| 2 | 1 | *i* | rotAziR2 | yes | -Inf | Inf | -25 | 25 | inherited | ° | $\theta_{R1}$ |
| 2 | 1 | *i* | rotEleR2 | yes | -Inf | Inf | -25 | 25 | inherited | ° | $\phi_{R1}$ |
| 2 | L1 | *e* | weight | yes | -1 | 1 | 0.001 | 0.999 | inherited | | $w_{bg}$ |

Note: In column *Parameter type*, *i* and *e* represent intrinsic and extrinsic parameters, respectively. In column *Model*, L1 represents layer 1. In column *initial value*, a tilde operator represents a median value. The function $pcaRot(x_k, y_k)$ outputs the angle between the x-axis and the first component of the localizations in the x-y plane. ‘inherited’ indicates that a value is inherited from its counterpart estimate in the last step. Other symbols have the
same meanings as in the main text. Abbreviations: LB/UB: lower/upper boundaries. Blanks indicate values not applicable.

**Supplementary Table 5.** Fitting settings used in this study for Nup96 in dual-color data.

| Step | Model | Parameter type | Internal parameter name | Free parameter | Relative |  | Absolute |  | Initial value | Unit | Symbol |
| --- | --- | --- | --- | --- | --- | --- | --- | --- | --- | --- | --- |
|  |  |  |  |  | LB | UB | LB | UB |  |  |  |
| 1 | 1 | *e* | x | yes | -50 | 50 | -150 | 150 | $\tilde{x}_k$ | nm | $x_0$ |
| 1 | 1 | *e* | y | yes | -50 | 50 | -150 | 150 | $\tilde{y}_k$ | nm | $y_0$ |
| 1 | 1 | *e* | z | yes | -100 | 20 | -300 | 300 | $\tilde{z}_k - 40$ | nm | $z_0$ |
| 1 | 1 | *e* | weight | no | | | | | 1 | | $w_1$ |
| 1 | 1 | *e* | xrot | yes | -30 | 30 | -inf | inf | 0 | ° | $\alpha$ |
| 1 | 1 | *e* | yrot | yes | -30 | 30 | -inf | inf | 0 | ° | $\beta$ |
| 1 | 1 | *e* | zrot | no | | | | | 0 | ° | $\gamma$ |
| 1 | 1 | *e* | xscale | no | | | | | 1 | | $s_x$ |
| 1 | 1 | *e* | yscale | no | | | | | 1 | | $s_y$ |
| 1 | 1 | *e* | zscale | no | | | | | 1 | | $s_z$ |
| 1 | 1 | *e* | variation | no | | | | | 0 | nm | $\epsilon$ |
| 1 | 2 | *e* | x | no | | | | | 0 | nm | $x_0$ |
| 1 | 2 | *e* | y | no | | | | | 0 | nm | $y_0$ |
| 1 | 2 | *e* | z | yes | -30 | 100 | 0 | 300 | 40 | nm | $z_0$ |
| 1 | 2 | *e* | xrot | no | | | | | 0 | ° | $\alpha$ |
| 1 | 2 | *e* | yrot | no | | | | | 0 | ° | $\beta$ |
| 1 | 2 | *e* | zrot | no | | | | | 0 | ° | $\gamma$ |
| 1 | 2 | *e* | variation | no | | | | | 0 | nm | $\epsilon$ |
| 1 | 2 | *e* | weight | no | | | | | 1 | | $w_1$ |
| 1 | 2 | *e* | xscale | no | | | | | 0 | | $s_x$ |
| 1 | 2 | *e* | yscale | no | | | | | 0 | | $s_y$ |
| 1 | 2 | *e* | zscale | no | | | | | 0 | | $s_z$ |
| 1 | L1 | *e* | weight | yes | -1 | 1 | 0.001 | 0.999 | 0 | | $w_{bg}$ |
| 2 | 1 | *e* | x | yes | -20 | 20 | -150 | 150 | $\hat{x}_0^{1,1} + \phi(\hat{z}_0^{1,2})_{[1]}$ | nm | $x_0$ |
| 2 | 1 | *e* | y | yes | -20 | 20 | -150 | 150 | $\hat{y}_0^{1,1} + \phi(\hat{z}_0^{1,2})_{[2]}$ | nm | $y_0$ |
| 2 | 1 | *e* | z | yes | -20 | 20 | -300 | 300 | $\hat{z}_0^{1,1} + \phi(\hat{z}_0^{1,2})_{[3]}$ | nm | $z_0$ |
| 2 | 1 | *e* | xrot | yes | -10 | 10 | -Inf | Inf | inherited | ° | $\alpha$ |
| 2 | 1 | *e* | yrot | yes | -10 | 10 | -Inf | Inf | inherited | ° | $\beta$ |
| 2 | 1 | *e* | zrot | yes | -180 | 180 | -inf | inf | [1 360] | ° | $r$ |
| 2 | 1 | *e* | variation | yes | -10 | 10 | 1 | 20 | 5 | nm | $\epsilon$ |
| 2 | 1 | *e* | weight | no | | | | | 1 | | $w_1$ |
| 2 | 1 | *e* | xscale | no | | | | | 1 | | $s_x$ |
| 2 | 1 | *e* | yscale | no | | | | | 1 | | $s_y$ |
| 2 | 1 | *e* | zscale | no | | | | | 1 | | $s_z$ |
| 2 | 1 | *i* | ringDistance | no | | | | | 50.2 | nm | $s$ |
| 2 | 1 | *i* | radius | no | | | | | 53.4 | nm | $r$ |
| 2 | 1 | *i* | cornerDegree | no | | | | | 12 | ° | $\psi$ |
| 2 | 1 | *i* | azimuthalShift | no | | | | | 8.8 | ° | $\theta$ |
| 2 | L1 | *e* | weight | yes | -1 | 1 | 0.001 | 0.999 | inherited | | $w_{bg}$ |

Note: In column *Parameter type*, *i* and *e* represent intrinsic and extrinsic parameters, respectively. In column *Model*, L1 represents layer 1. In column *initial value*, a tilde operator represents a median value, a hat operator indicates a parameter estimate, and superscripts represent steps and models (e.g., $\hat{z}_0^{1,2}$ means the estimate of $z_0$ from model 2 in step 1). The function $\tilde{z}_{[k]}' = \phi(z)_{[k]}$ outputs the $k^{\text{th}}$ element of the vector $\tilde{z}'$ derived from $\tilde{z} = [0 \ 0 \ z]$ rotated by the rotation matrix constructed using $\{\hat{\alpha}^{1,1}, \hat{\beta}^{1,1}, \hat{\gamma}^{1,1}\}$ as the angles $\{\alpha, \beta, \gamma\}$ in the equation \(11\). ‘inherited’ indicates that a value is inherited from its counterpart estimate in the last step. Other symbols have the same meanings as in the main text. Abbreviations: LB/UB: lower/upper boundaries. Blanks indicate values not applicable.

**Supplementary Table 6.**
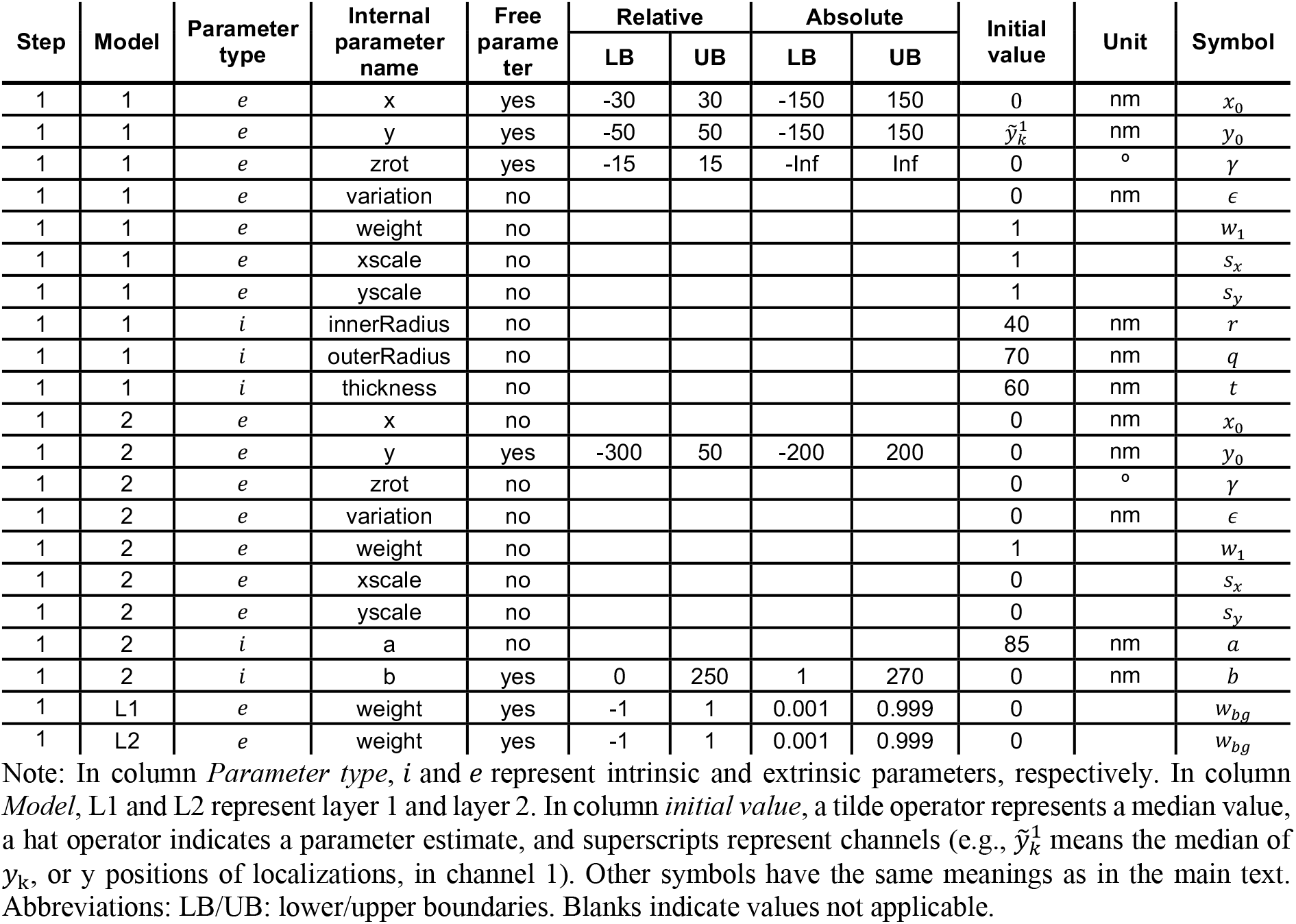
Fitting settings used in this study for endocytic sites in yeast.

| Step | Model | Parameter type | Internal parameter name | Free parameter | Relative |  | Absolute |  | Initial value | Unit | Symbol |
| --- | --- | --- | --- | --- | --- | --- | --- | --- | --- | --- | --- |
|  |  |  |  |  | LB | UB | LB | UB |  |  |  |
| 1 | 1 | *e* | x | yes | -30 | 30 | -150 | 150 | 0 | nm | $x_0$ |
| 1 | 1 | *e* | y | yes | -50 | 50 | -150 | 150 | $\tilde{y}_k^1$ | nm | $y_0$ |
| 1 | 1 | *e* | zrot | yes | -15 | 15 | -Inf | Inf | 0 | ° | $\gamma$ |
| 1 | 1 | *e* | variation | no | | | | | 0 | nm | $\epsilon$ |
| 1 | 1 | *e* | weight | no | | | | | 1 | | $w_1$ |
| 1 | 1 | *e* | xscale | no | | | | | 1 | | $s_x$ |
| 1 | 1 | *e* | yscale | no | | | | | 1 | | $s_y$ |
| 1 | 1 | *i* | innerRadius | no | | | | | 40 | nm | $r$ |
| 1 | 1 | *i* | outerRadius | no | | | | | 70 | nm | $q$ |
| 1 | 1 | *i* | thickness | no | | | | | 60 | nm | $t$ |
| 1 | 2 | *e* | x | no | | | | | 0 | nm | $x_0$ |
| 1 | 2 | *e* | y | yes | -300 | 50 | -200 | 200 | 0 | nm | $y_0$ |
| 1 | 2 | *e* | zrot | no | | | | | 0 | ° | $\gamma$ |
| 1 | 2 | *e* | variation | no | | | | | 0 | nm | $\epsilon$ |
| 1 | 2 | *e* | weight | no | | | | | 1 | | $w_1$ |
| 1 | 2 | *e* | xscale | no | | | | | 0 | | $s_x$ |
| 1 | 2 | *e* | yscale | no | | | | | 0 | | $s_y$ |
| 1 | 2 | *i* | a | no | | | | | 85 | nm | $a$ |
| 1 | 2 | *i* | b | yes | 0 | 250 | 1 | 270 | 0 | nm | $b$ |
| 1 | L1 | *e* | weight | yes | -1 | 1 | 0.001 | 0.999 | 0 | | $w_{bg}$ |
| 1 | L2 | *e* | weight | yes | -1 | 1 | 0.001 | 0.999 | 0 | | $w_{bg}$ |

Note: In column *Parameter type*, *i* and *e* represent intrinsic and extrinsic parameters, respectively. In column *Model*, L1 and L2 represent layer 1 and layer 2. In column *initial value*, a tilde operator represents a median value, a hat operator indicates a parameter estimate, and superscripts represent channels (e.g., $\tilde{y}_k^1$ means the median of $y_k$ , or y positions of localizations, in channel 1). Other symbols have the same meanings as in the main text. Abbreviations: LB/UB: lower/upper boundaries. Blanks indicate values not applicable.

